# The dynamical Matryoshka model: 2. Modeling of local lipid dynamics at the sub-nanosecond timescale in phospholipid membranes

**DOI:** 10.1101/2022.03.30.486370

**Authors:** Aline Cisse, Tatsuhito Matsuo, Marie Plazanet, Francesca Natali, Michael Marek Koza, Jacques Ollivier, Dominique J. Bicout, Judith Peters

**Affiliations:** Univ. Grenoble Alpes, CNRS, LiPhy, 38000 Grenoble, France; Institut Laue-Langevin, 71 avenue des Martyrs, CS 20156, 38042 Grenoble Cedex 9, France; CNR-IOM and INSIDE@ILL, c/o OGG, 38042 Grenoble Cedex 9, France; Univ. Grenoble Alpes, CNRS, Grenoble INP, VetAgro Sup, TIMC, 38000 Grenoble, France; Institute for Quantum Life Science, National Institutes for Quantum Science and Technology, 2-4 Shirakata, Tokai, Ibaraki, 319-1106, Japan; Institut Universitaire de France

**Keywords:** Lipid membranes, neutron scattering, molecular dynamics, modeling

## Abstract

Biological membranes are generally formed by lipids and proteins. Often, the membrane properties are studied through model membranes formed by phospholipids only. They are molecules composed by a hydrophilic head group and hydrophobic tails, which can present a panoply of various motions, including small localized movements of a few atoms up to the diffusion of the whole lipid or collective motions of many of them. In the past, efforts were made to measure these motions experimentally by incoherent neutron scattering and to quantify them, but with upcoming modern neutron sources and instruments, such models can now be improved. In the present work, we expose a quantitative and exhaustive study of lipid dynamics on DMPC and DMPG membranes, using the Matryoshka model recently developed by our group. The model is confronted here to experimental data collected on two different membrane samples, at three temperatures and two instruments. Despite such complexity, the model describes reliably the data and permits to extract a series of parameters. The results compare also very well to other values found in the literature.

Figure 1:
Graphical abstract.

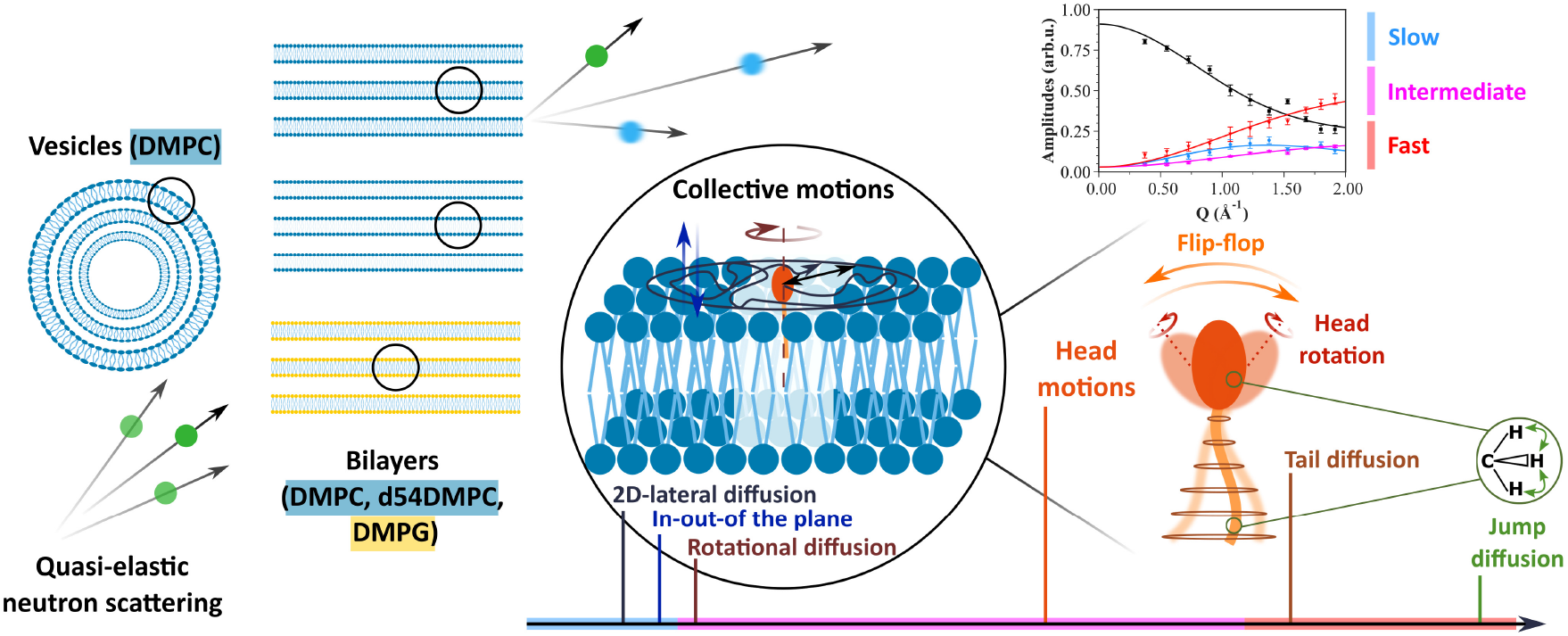

**Highlights:**

- The Matryoshka model brings a new general description of local lipid dynamics.
- Phospholipid membranes on various conditions are compared in this novel framework.
- Effects of main phase transition, membrane geometry or motion direction are probed.
- Despite high number of parameters, overfitting is avoided by a global fit strategy.

## 1 Introduction

Lipid membranes are at the basis of cell organization. Enabling a clear separation between cell constituents and the external solvent, they also permit to partition the different components inside the cell, delineating for example the genetic material in a nucleus for eukaryotic cells [1]. Depending on their composition, their association with other molecules, like cholesterol or membrane proteins, they act as borders, and filter what enters or leaves the cell.

Therefore, the study of the structure of lipid membranes is of primary importance to link it to its functionality. Membranes can directly be visualized by cryo-electron microscopy (cryo-EM) [2] [3] or atomic force microscopy (AFM) [4] [5]. Neutron scattering techniques have the advantage to be non-invasive and to permit investigating structure and dynamics at the length and time scales appropriate for lipids and membranes. Scattering methods, including small-angle scattering (SAS) [6] [7], diffraction [8] [9] [10] or reflectometry [11], enable to access high-resolution information, like bilayers’ spacing, radii of vesicles and even structural features from the lipids, such as the head group size or volume [10] [12] [13]. Studies on the effect of lipid composition, addition of molecules, assembly with membrane proteins, under different conditions as hydration [14], temperature [8] or pressure [15], are numerous.

However, membrane functionality is not only led by the structure, but also by the dynamics. For example, it was shown for bacteriorhodopsin in purple membranes [16] that the hydration level as well as the dynamics of the membrane and the protein are related to the functionality of the whole system. Photosynthetic membranes also present strong correlations between functionality and dynamics [17] [18]. Among the techniques that can be used, we can cite fluorescence techniques [19], nuclear magnetic resonance (NMR) [20] [21], dynamic light scattering (DLS) [22], neutron scattering including spin-echo (NSE) for long time-scales [23] [24], and elastic and quasi-elastic incoherent neutron scattering (EINS and QENS) at shorter times [14] [25]. The properties of the membranes are governed by their thermodynamic characteristics, which in turn can be measured by differential scanning calorimetry (DSC) [8].

Incoherent neutron scattering is well adapted to probe the sub-nanosecond dynamics of lipids in membranes. As neutrons are sensitive to hydrogen atoms (H), which constitute around 50 % of biological samples, they enable to see various molecular motions without sample damage. Neutron instruments permit to study all lipid membrane geometries, multilamellar bilayers (MLBs) or multilamellar vesicles (MLVs) as well as non-lamellar structures, with the possibility of focusing on in-plane or out-of-plane motions for the MLBs. Finally, neutrons are also sensitive to isotopic substitution : concerning incoherent scattering, the scattering cross section of hydrogen is around 40 times higher compared to deuterium atoms [26]. As a consequence, it is possible to spotlight certain parts of a sample by deuterating the other parts. In the case of lipids, it is particularly interesting to deuterate the tail(s) to have a focus on the head dynamics or vice versa.

Though, QENS spectra, giving experimentally access to the dynamic structure factor 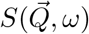, need a model to interpret the data and describe precisely the lipid dynamics. The model presented by Pfeiffer et al. in 1989 [25] was one of the first to describe the main types of motions occurring in lipid membranes, and to analyze QENS spectra accordingly. It included up to six possible movements in lipid membranes: (1) chain defect motions of the tails, (2) rotational diffusion along the long axis of a lipid molecule, (3) lateral diffusion in the plane of the membrane, (4) rotations and head-flip-flop motions, (5) vertical out-of-plane motions and (6) collective undulations. Having established this, Elastic Incoherent Structure Factors (EISF) from the spectrometers IN10 (time window about 1 ns) and IN5 (time window about 20 ps) at the Institut Laue Langevin (ILL) were fitted successfully to extract diffusion constants or the distance of protons from the rotational axis [25]. Other studies following a similar approach can be found in [27], [28], or [29].

However, the neutron flux was much lower in that time and so the statistics of experimental data worse, preventing the use of too many free parameters without overfitting the data. The rise of available neutron flux and data quality emerging from new spectrometers called for novel general models for lipid dynamics, which was first addressed in the work of Wanderlingh et al. [30] [31]. It supposes that within a very good approximation, motions in lipid membranes can be considered as dynamically independent and therefore separated in three time domains. Fast motions, at a sub-ps scale, include H motions with respect to the acyl carbon atoms and are described as an uniaxial rotational diffusion. At around 6 - 7 ps, intermediate motions comprise conformational dynamics in the lipid chains and rotational diffusion of the methyl groups. For these motions, the authors subdivided the lipid molecule into different parts, called “beads”, along the head groups and the tails and formulated their model for these beads. Finally, slow motions (40 - 350 ps) are described as translational diffusion of the whole phospholipid in-plane and out-of-plane. Such description was very successful when applied to the EISF.

Gupta et al. [32] developed a first model in 2018 to predict the dynamics of phospholipid membranes extracted from DLS and NSE measurements. The analysis of NSE data was based on the Zilman-Granek model combined with translational center-of-mass diffusion and further on a cumulant expansion to extract the mean square displacements. The large time window from 3 to 180 ns of NSE permitted then to identify three different power laws in time and the associated dynamics. More recently, Gupta et al. [24] proposed a new model for shorter and long time dynamics as probed by QENS (t < 5 ns) and spin-echo spectroscopy (t > 100 ns) on liposomes. Local motions at the shorter time scale include tail motions confined within a volume of cylindrical symmetry. Head group motions are taken into account only as a constant background as the head contains much less H atoms than the tails (typically a proportion 1:4). Long time motions are described as height-height correlations, thickness fluctuations and translational diffusion of the liposomes. The intermediate scattering functions are well described by this model as well as mean square displacements at the longer time scales. The model is therefore complementary to our approach, which applies to local motions and shorter time scales.

In the present investigation, we go beyond the cited studies, taking advantage on one hand of recent QENS results from the spectrometers IN6 and IN5 at ILL, and on the other hand of structural parameters known experimentally as initial values for the fits. However, one should have in mind that the parameters determined here are the values as seen from molecular motions and are not necessarily matching the static ones. The so-called Matryoshka model (the name is inspired from the nested Russian dolls to account for the hierarchy of motions) was first introduced by D.J. Bicout et al. [33] and validated against data from phospholipid bilayers of 1,2-dimyristoyl-*sn*-glycero-3-phosphocholine (DMPC) at three different temperatures. The model includes the main local motions as 2D-diffusion of the tails in a cylinder, jump-diffusive movements [34] of both tails and heads and head-flip-flop plus rotational diffusion in the head group as well as collective motions, all supposed to be independent of each other. In addition, across the three considered timescales and the elastic line, the amplitudes share the same parameters. In that way, the Matryoshka model can be applied in a global way to all the amplitudes, which constrains the fit, reduces overfitting, and increases the precision of the calculations.

The present paper exposes a quantitative and exhaustive study of lipid dynamics on DMPC and 1,2-dimyristoyl-*sn*-glycero-3-phospho-(1’-rac-glycerol) (DMPG) membranes, using the Matryoshka model and taking into account known features such as the main phase transition, membrane geometry, direction of motions, or lipid composition. The Matryoshka model will be first introduced, along with its main hypotheses. Then, details on the samples, neutron scattering experiments and subsequent analyses will be given. Finally, results will be presented and discussed.

## 2 The Matryoshka model for lipid dynamics

### 2.1 Main hypotheses

As shown in Figure 2a, in a simplified way a lipid is assumed here as constituted of two bodies: a head group, containing a fraction *z* of H atoms, and an effective tail, containing a proportion (1 − *z*) of the total number of H atoms. As lipid molecules, like DMPC or DMPG, are usually made of more than one tail, the effective tail in Figure 2a is used to represent and describe motions of all H atoms in tails regardless their belonging. However, neutron scattering does not allow to distinguish if a H atom belongs to one or the other tail, as only averaged motions are measured. Moreover, data fitting and the resulting parameters show very consistent outcomes, which strengthen our approach.

**Figure 2:**
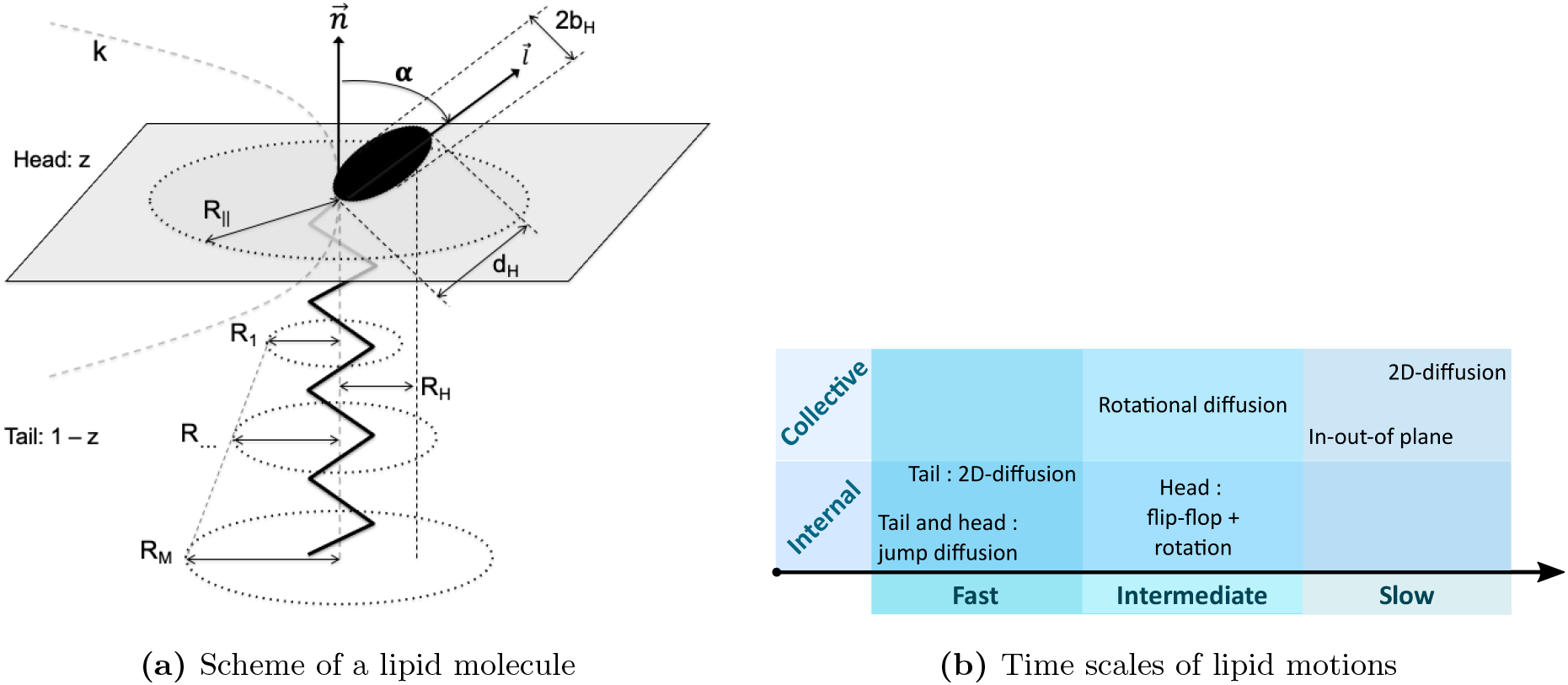
Main elements of the model. 2a : representation of a lipid molecule in the membrane and associated parameters (reproduced from [33]). 2b : hierarchy of the motions considered in the Matryoshka model.

The Matryoshka model assumes six types of lipid motions, whatever the samples studied or the instrumental resolution of the experiment. These motions are ranked in terms of their time scales (noted as fast, intermediate or slow, similar to [30]), and are subdivided into collective (concerning a motion of the whole lipid) or internal movements (only parts of the lipid; methylene and methyl groups, head or tail), as described in Figure 2b.

The slowest motions consider only the whole lipid, and are happening in two opposite directions with regard to the membrane :

- A lipid can freely 2D-diffuse within the membrane plane, and more particularly in an average cylinder of radius *R*_∥_, as schematized in Figure 2a. *R*_∥_ will be referred as the lateral diffusion radius in the following.
- The same lipid can also oscillate in the out-of-plane direction, moving normal to the membrane plane. This motion is characterized by a force constant *k_force_* : when *k_force_* is high, it means the membrane is rigid, and the lipid moves only little in the out-of-the plane direction, whereas a small *k_force_* means a higher flexibility.

The motions said to be intermediate are a mixture of internal and collective dynamics :

- The whole lipid can rotate around its normal axis. The rotational diffusion expression contains the half-height of the head group, *R*_H_ (see Table S1).
- Inside the lipid, the head group can perform a head-flip-flop motion between the angle *α* and −*α* with respect to the normal axis. In addition to this head-flip-flop motion, the head with radius *b*_H_ can rotate around its own axis (see Table S1).

Finally, the fast motions are only internal and concern different parts of the lipid :

- The lipid tail can 2D-diffuse around the normal axis. But the extension of that motion will change depending on the H position in the tail : close to the head, the motions are seen more restricted, around a radius *R*_1_, whereas far from the head, the motions can extend until 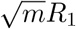, with *m* the index of the methylene and methyl group position on the tail, and *M* the total number of these groups.
- In addition, the jump-diffusion of the H atoms inside the methylene and methyl groups has to be taken into account. Jump-diffusion assumes to occur via infinitely small, elementary jumps characterized by a negligible jump time during which the particle diffuses, and the residence time *τ*, i.e. the time a particle spends in a given position [34]. The methylene and methyl groups can be found in the tail, but also in the head. The involved parameters are the distance H-C-H *d* between the two-sites concerned by the jump-diffusion, but also the probability *ϕ* of jump events (see Table S1).

Assuming faster motions for tails than for heads can appear counter-intuitive, as they are bigger and buried inside the membrane. However, in addition to be supported by the following results, other studies showed that tails’ motions set in at lower temperature than those of the head groups, and seem even to drive head motions [35]. Assuming a shorter time scale and fastest motions for tails appears then to be a reasonable hypothesis.

## 3 Materials and Methods

### 3.1 Samples

DMPC (1,2-dimyristoyl-*sn*-glycero-3-phosphocholine) and DMPG (1,2-dimyristoyl-*sn*-glycero-3-phospho-(1’-rac-glycerol) (sodium salt)), represented in Fig. 3, were purchased from Lipoid (Ludwigshafen Germany) or from Avanti Polar Lipids (Alabaster, USA) and used without further purification.

**Figure 3:**
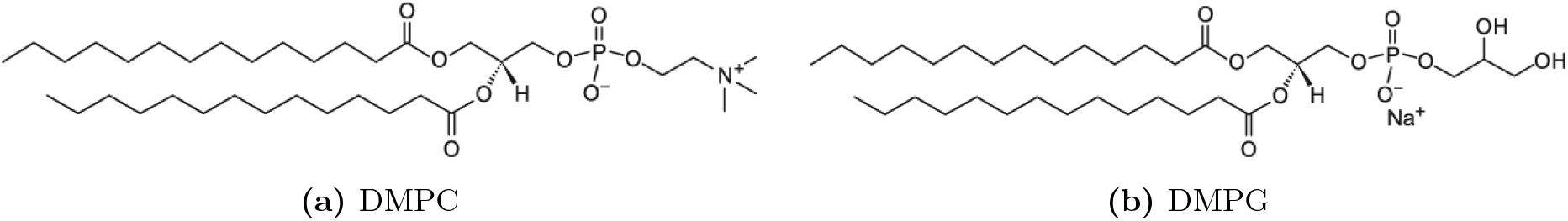
Molecular representations of both lipid samples. (Reproduced from avantilipids.com).

**Figure 4:**
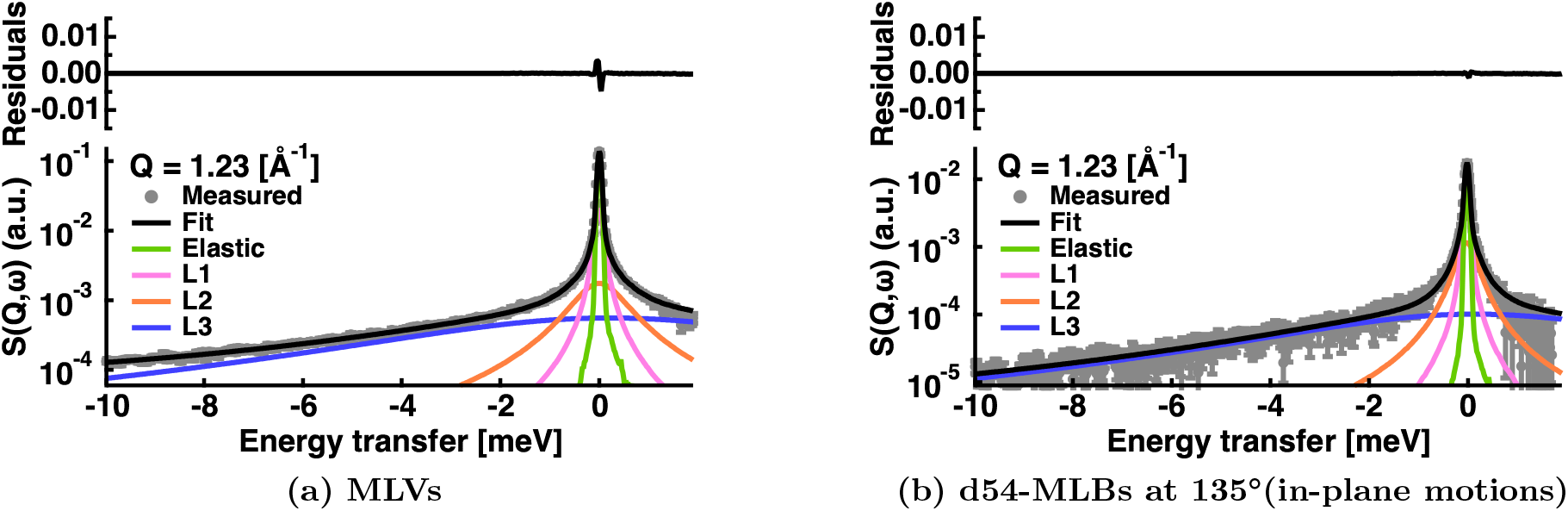
Example of S(*Q* = 1.23 Å^−1^, *ω*) data fitting for two samples measured on IN6 at T = 283 K. The grey circles are the data points. The total fit is represented by the black line. The green line corresponds to the elastic peak convoluted with the resolution function, which is directly given by the Vanadium measurements at *Q* = 1.23 Å^−1^. The magenta, orange and blue curves are respectively the Lorenzian functions convoluted by 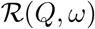 for slow, intermediate and fast motions. Residuals are showed at the top of each figure.

Different sample geometries were investigated on two neutron spectrometers : MLVs, MLBs and MLBs, whose lipid tails were deuterated (d54-MLBs) so that only the heads were visible by neutrons. The samples used on the two instruments were different, but prepared following the same protocol (see [36]).

Shortly, for MLVs, about 100 mg of lipid powder were placed in a flat sample holder and hydrated in a desiccator from pure D_2_O for two days at 40 *°*C. Additional heavy water was added to achieve a sample with an excess of water [37]. DMPC and DMPG oriented MLBs, with a full width half-maximum of a neutron rocking curve of the first Bragg peak typically between 1 and 3°, were prepared on Si wafers and hydrated with heavy water. We used the “rock and roll” method following a protocol described by Tristram-Nagle and co-workers [38] in which DMPC powder was deposited on a Si(111) wafer of dimensions 30 × 40 × 0.38 mm^3^ by evaporating from a trifluoroethanol: chloroform mixture (2:1, v/v). After deposition, the wafer was dried over silica gel for 2 days in a desiccator. The sample was rehydrated from pure D_2_O at 40 *°*C to achieve a high hydration level (we used 27 % weight of water on IN6 against 10 % on IN5). One wafer contained a total amount of ≈ 35 mg of lipids.

Both MLVs or MLBs were placed in slab-shaped aluminum sample holders, gold-coated to avoid sample contamination. Sample cells were sealed using indium wire and the weight of the sample was monitored before and after the experiment, with no change observed indicating a stable level of hydration.

### 3.2 Neutron scattering experiments

DMPC samples with the three geometries (MLVs, MLBs, d54-MLBs) were measured on the IN6 time-of-flight spectrometer from ILL (Grenoble, France), with a wavelength of 5.1 Å, corresponding to an energy resolution of 75 μeV [39]. At this resolution, motions up to around 10 ps are accessible, and the attainable Q-range is of [0.37; 2.02] Å^−1^. Quasi-elastic neutron scattering (QENS) scans were performed at three different temperatures, 283 K, 311 K and 340 K, to probe the dynamics before and after the main phase transition of the lipids (at 297 K for DMPC and 296 K for DMPG). For MLBs, to access in-plane or out-of-plane motions, the sample holder was oriented at 135°, respectively 45°, with respect to the beam (see [40] for example).

To have a comparison with another sample, as well as a different instrumental resolution, data from an experiment performed on the IN5 time-of-flight spectrometer from ILL (Grenoble, France) were analyzed in addition [41]. Here, DMPC and DMPG samples in MLB geometry were scanned at 280 K and 295 K, with a wavelength of 6 Å. This configuration corresponds to an energy resolution of about 45 μeV, and an observable time scale of 15 ps. The accessible Q-range was of [0.18; 1.82] Å^−1^.

In both experiments, an empty cell with and without wafers, as well as Vanadium, were measured for correction and normalization purposes. In order to avoid multiple scattering, the sample thickness was calculated to give a transmission of about ~90 %.

### 3.3 QENS analysis

Raw data were first corrected for the empty cell and the contribution of six wafers, using the Large Array Manipulation Program (LAMP) [42].

The resulting 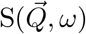 spectra were subsequently analyzed in the range of −10 meV ≤ ΔE ≤ 2 meV using IGOR Pro software (WaveMetrics, Lake Oswego, OR, USA). The general model used for fitting the spectra [43] was the following :

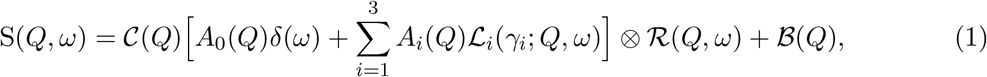

with 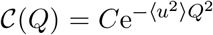, the Debye-Waller factor, *A*_0_ the elastic incoherent structure factor (EISF), *A_i_* and *γ_i_* the respective amplitudes and half-widths at half-maximum of the Lorentzian functions 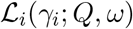. In most of QENS studies, the amplitudes *A_i_* obtained through this “model-free” approach are ignored and not further exploited during data analysis. However, they allow to shed light on various structural-dynamical aspects of the samples, as shown in the following. Moreover, for each sample and temperature, the EISF *A*_0_ and the three amplitudes *A_i_* are fitted globally, which reduces the risk of overfitting. For all these reasons, here the amplitudes *A*_0_ as well as *A_i_* were subsequently analysed with the Matryoshka model, allowing to extract a series of parameters. In a follow-up study [44], the *γ_i_* are investigated in more details using either classical models or the same Matryoshka model; we will refer to it by “the linewidths analysis” in the text. 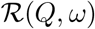 refers to the resolution function, and corresponds to the Vanadium measurements, directly included in the analysis. ⊗ designates a convolution. Finally 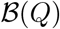 is a flat background, that can comprise the instrumental contribution, or fast vibrational motions which are too flat to be analysed separately. Fig. 3 shows two examples of experimental data at *Q* = 1.23 Å^−1^ together with fitted curves according to Equation 1.

Similarly to the QENS analysis in [30], three Lorentzian functions were used for fitting, accounting for three diffusional processes with distinct relaxation times. Each process is assumed to correspond to a certain lipid motion from Table S1 (Supplementary Material), following the hierarchy of motions presented in Fig. 2b. The Table 1 summarizes which motions reside within each timescale.

**Table 1:** Lipid motions within each timescale.

| Timescale dynamics | Dynamical processes |
| --- | --- |
| Slow | in-out of the plane $\otimes$ 2D lateral diffusion |
| Intermediate | head motions $\otimes$ rotational diffusion |
| Fast | tail motions (2D-diffusion $\otimes$ jump-diffusion)<br>+ jump-diffusion of H in head |

### 3.4 The theoretical fit model

The association of lipid motions to a corresponding timescale, presented in Table 1, enables to write the theoretical EISF and the amplitude of each Lorentzian function of Eq. (1) (see Table 2).

**Table 2:** Fit functions used for the amplitudes. *z* is the proportion of hydrogen atoms in the head. *m* refers to the mobile fraction of H atoms. *ε*_0_ is a factor accounting for the immobile fraction of H atoms, but also for multiple scattering effects, which can become visible at low-Q range [45, 46]. A homogeneous distribution over length scales of the errors is assumed, and thus *ε*_0_(*Q*) = *ε*_0_ (see [33]).

| Amplitudes | Theoretical function |
| --- | --- |
| $A_0$ (EISF) | $\varepsilon_0 + m \left[ [(1-z)A_{tail}A_{jd} + zA_{head}A_{jd}] A_{rot} A_{in-out} A_{2d} \right]$ |
| $A_1$ (slow) | $\frac{1-m-\varepsilon_0}{3} + m \left[ [(1-z)A_{tail}A_{jd} + zA_{head}A_{jd}] A_{rot} (1 - A_{in-out} A_{2d}) \right]$ |
| $A_2$ (intermediate) | $\frac{1-m-\varepsilon_0}{3} + m \left[ zA_{jd}(1 - A_{head}A_{rot}) + (1-z)A_{tail}A_{jd}(1 - A_{rot}) \right]$ |
| $A_3$ (fast) | $\frac{1-m-\varepsilon_0}{3} + m \left[ z(1 - A_{jd}) + (1-z)(1 - A_{tail}A_{jd}) \right]$ |

Thus, the application of the theoretical model for various motions of the lipids presented in Section 2 is performed on the EISF and the amplitudes. The latter are corresponding to the areas under the Lorenzian curves retrieved from the previous QENS analysis. They are fitted with the functions of Table 2, using the package *lmfit* from Python [47], with the implemented Levenberg-Marquardt and Nelder-Mead algorithms [48, 49], following a three-step procedure:

1. A fit is performed using the Nelder-Mead algorithm to get first estimations of the parameters. Contrary to the common Levenberg-Marquardt optimization, it is a direct search algorithm which does not require the calculations of derivatives. Several tests we ran indicated that the Nelder-Mead option was less influenced by the initial parameters, while staying quite efficient in terms of time and consistency of the values.
2. To have an estimation of the error bars, the Levenberg-Marquardt algorithm is then used, using the previous results from the Nelder-Mead fit.
3. Finally, to assess the effect of each parameter on the global fit, all parameters except one are fixed, and the Levenberg-Marquardt algorithm is applied another time. The returned value and error bar are saved, and the process is repeated for each parameter (except for the parameters which are known from experiments and fixed. They are summarized in Table 3).

**Table 3:** Parameters known from experiments.

| Parameter | Value | Literature |
| --- | --- | --- |
| $M$ | 14 | Sample datasheet [50] |
| $z$ | 0.25 (DMPC), 0.18 (DMPG) | Sample datasheet [50] |
| $\alpha$ ($^\circ$) | $32.3 \pm 0.6$ (DMPC), 30 (DMPG) | Diffraction values [38], [51] |
| $d$ ($\text{\AA}$) | $2.2 \pm 0.1$ | Average C-H distance (CCCBDB from NIST [52]) |

The fitting procedure relies at each step on the simultaneous fitting of the four areas with their corresponding fit function (gathered in Table 2), through a set of shared parameters (see Table S1 for the list of parameters). In such a way, statistics are improved by the use of more data points, but it also constrains the fit, and limits the combination of parameters that could lead to a good fit.

The quality of the fits is given through the reduced chi-square value of all shown fit examples (see Figures 5, 6 and Appendix B Figures 1, 2, 3). In general, overfitting leads to the fitted curves that unnaturally pass through all the data points within their error bars. While our fits in Figures 5 and 6 reproduce the experimental profiles quite well as a whole, they do not pass through several data points, implying that the number of variables in the model is less than that required for overfitting. We also refer to Appendix E, which explains that even slight changes in the model provoke a mismatch of the fits. All extracted parameters are given with their error bars in the Figures 7 and 8, in the tables in Appendix D (Tables 2–7), which also prove that the fits work remarkably well.

**Figure 5:**
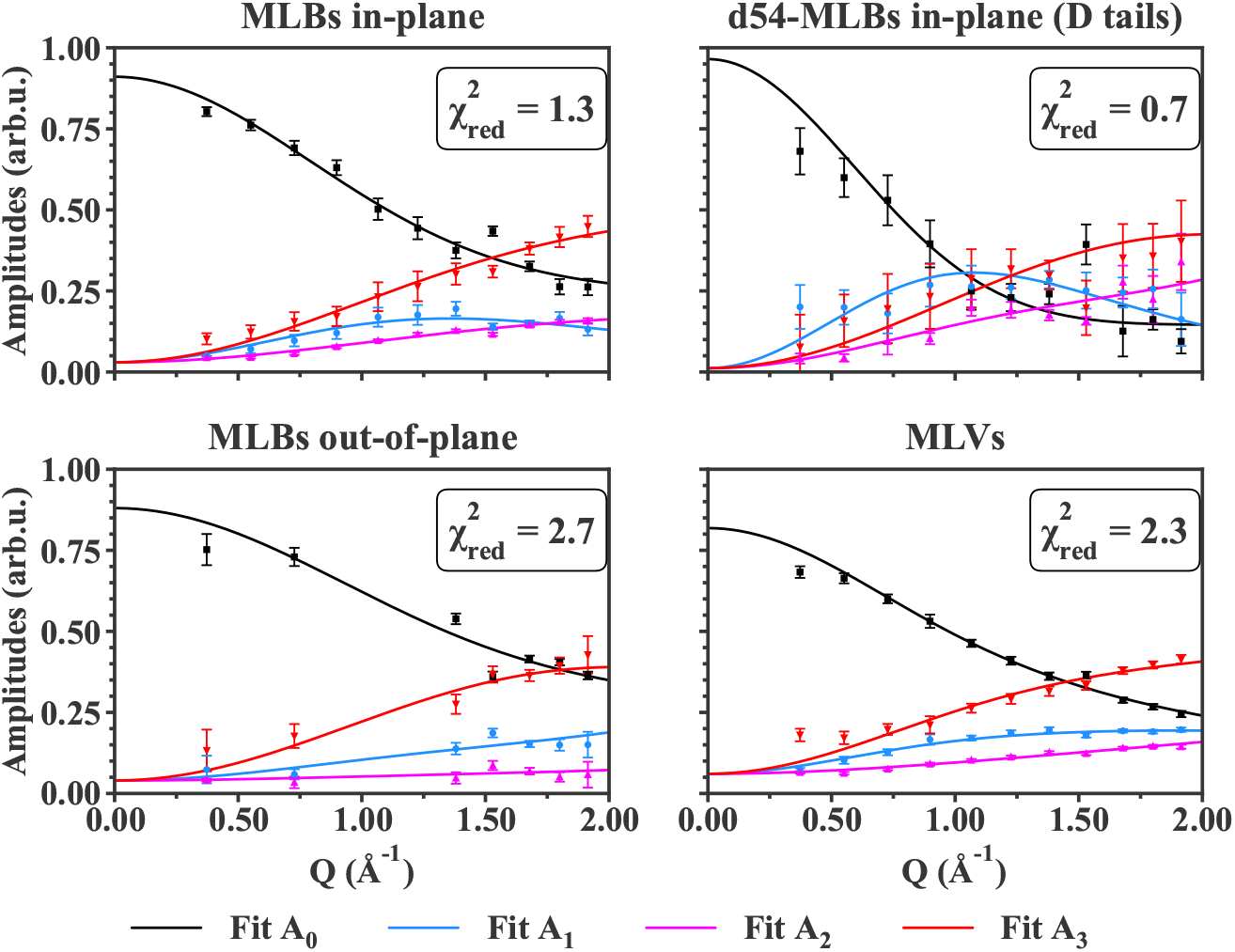
Fit curves against data points for IN6 data at T = 283 K for all samples measured. The black points and line correspond to the EISF (*A*_0_) data points and fit curve. In the same manner, the blue points and line represent *A*_1_, the amplitude which coincides with the slowest motions within the instrumental resolution. The magenta points and line are linked to *A*_2_, whose motions timescales are said intermediate. Finally, *A*_3_, linked to the fastest motions, is represented by red points and line. 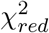 indicates the reduced chi-square value of the fit. Error bars are within symbols if not shown.

**Figure 6:**
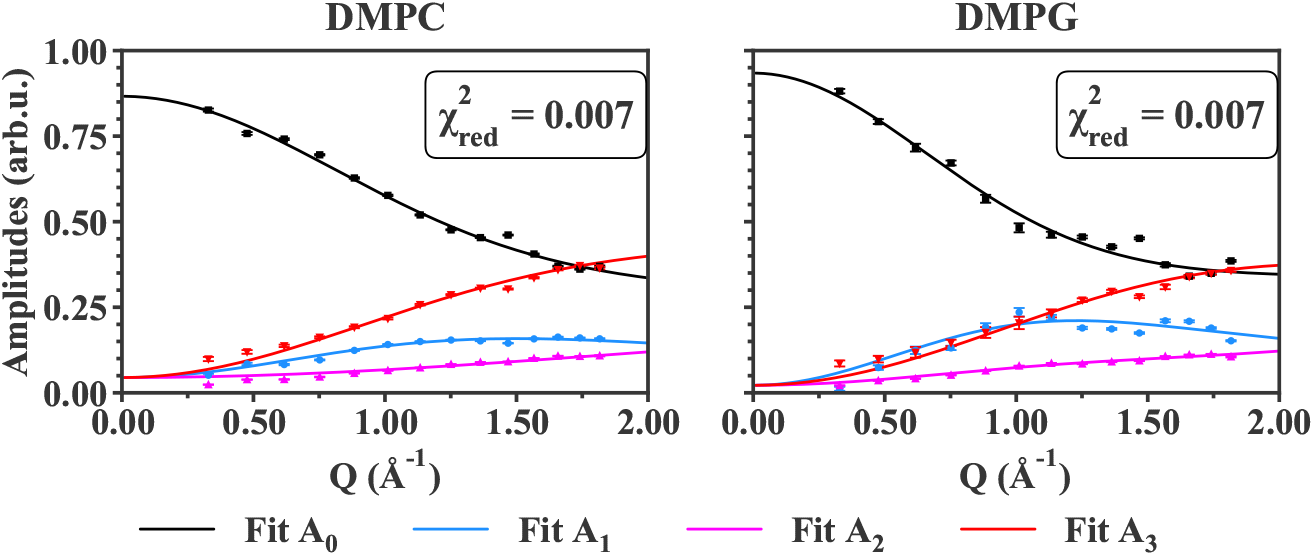
Fit curves against data points for IN5 data at T = 280 K for all samples measured. The legend is the same as in the previous Figure 5 for IN6 data. Error bars are within symbols if not shown.

**Figure 7:**
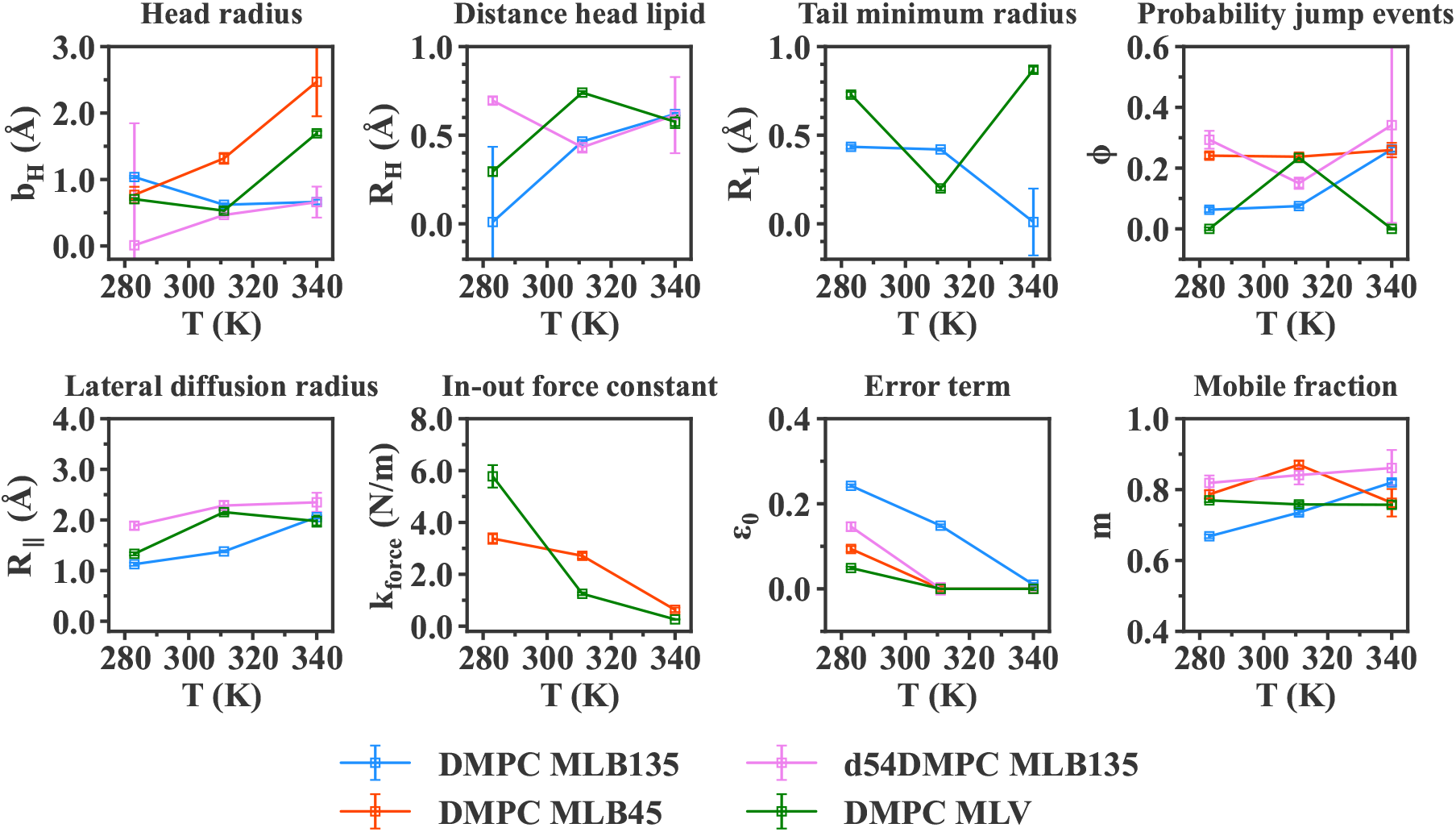
Comparison of each parameter between IN6 DMPC samples. In blue is represented MLB sample measured at 135° (in-plane motions). The MLB sample measured at 45° (out-ofplane motions) is depicted in orange. The tails-deuterated sample, d54-MLB, is is pink. Finally MLV sample is shown in green. Missing values occur when the parameter is not present in the model for a specific sample (for example, the parameter *R*_∥_ prevails only for in-plane motions). Error bars are within symbols if not shown.

**Figure 8:**
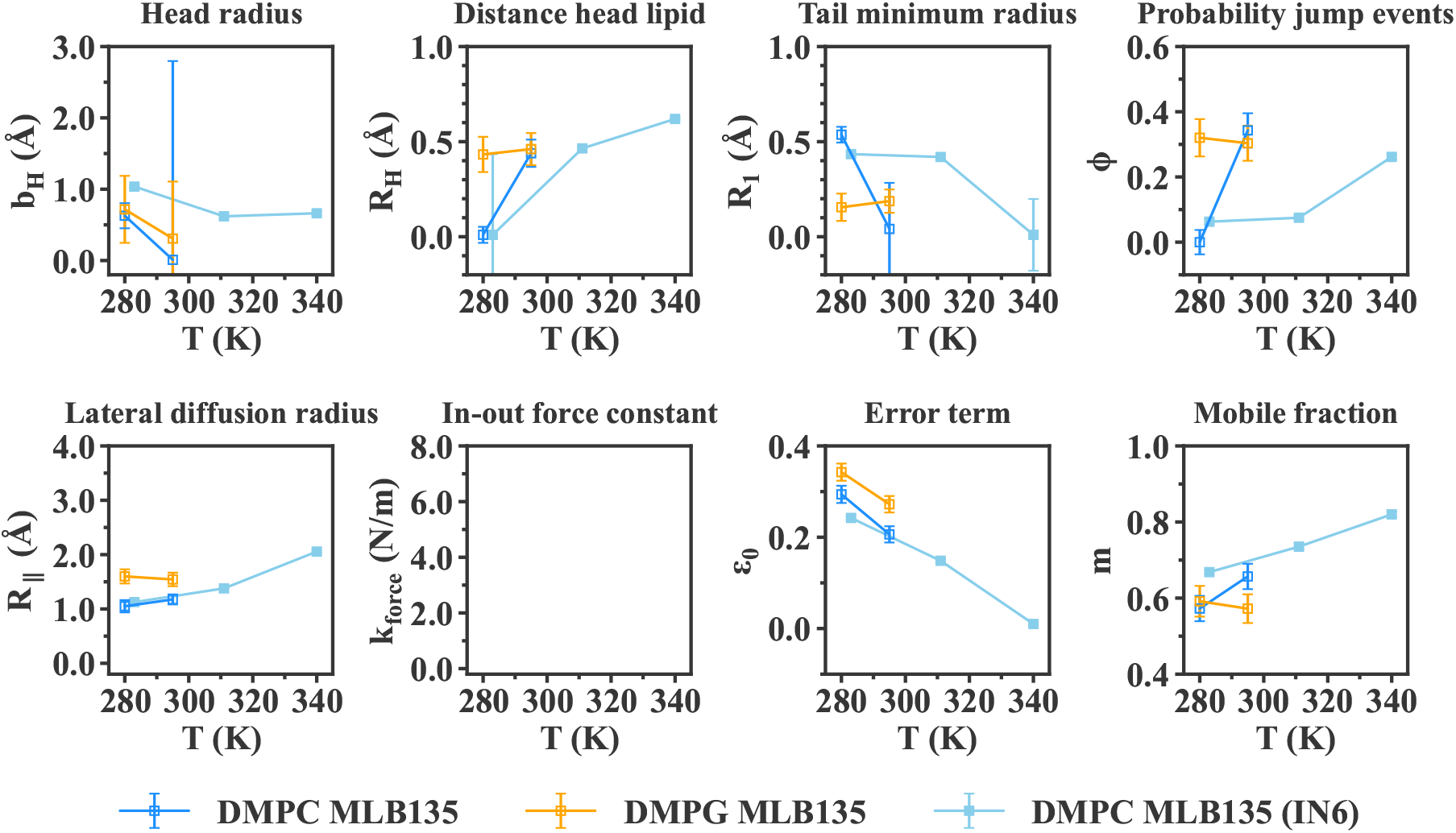
Comparison of each parameter between IN5 samples, both MLB measured at 135° (in-plane motions). In blue is represented DMPC MLB sample, whereas DMPG is represented in yellow. For comparison, the DMPC MLB sample measured on IN6 is shown in filled light blue squares. Missing values occur when the parameter is not present in the model for a specific sample. Error bars are within symbols if not shown.

Finally, the Matryoshka model directly differentiates the directions of motions in the theoretical expressions of the amplitudes, allowing to write different functions for in-plane or out-of-plane motions, and thus for MLBs measured at 135° or 45°(see [33], Table 4). In the case of MLVs, an average of the amplitudes at four different directions (in-plane, out-of-plane, and two intermediate angles) is computed to lead to the total amplitudes *A_i_*. Choosing four directions for averaging was proved to be sufficient enough to describe the data, while keeping a reasonable computational time.

Some parameters were fixed according to experimental values which can be found in the literature. We summarize these parameters in the following table, with the corresponding references :

## 4 Results

The amplitudes *A_i_*, including the elastic incoherent structure factor (EISF, i=0) and the quasi-elastic incoherent structure factors (QISF, i=1,2,3) retrieved from the QENS analysis, were fitted as explained in section 3. The comparison between the data points and the fit curves are shown for all samples, measured on both IN6 and IN5 instruments, for T = 283 K and T = 280 K, respectively, in Figures 5 and 6. The higher temperatures studied are displayed in the Supplementary Material, in Figures S1, S2 and S3.

For all the samples measured, whatever the lipid geometry, type of motions or lipid composition, the fit curves approach very well the data points. The reduced chi-square values (see in Figure 5) are around 1 to 2, indicating a good fit quality. Concerning the IN5 data, the error bars of the amplitudes are ten times smaller than on IN6. Numerically, error weighted fits were then too restricted. Even if weighted and non-weighted fits led to similar results, non-weighting remained the most stable option, and was preferred in the case of IN5 data. For that reason, the reduced chi-square values presented in Figure 6 have not the same definition (as it does not account for the weights) and correspond to the averaged squared residuals, which are then close to zero. However, the curves match again the data points, and clearly follow the experimental trends.

In all cases, the Matryoshka model describes well the decrease of *A*_0_, the EISF, and the increase of the QISF, *A*_1,2,3_, with increasing Q-values. Notably, the behaviour of *A*_1_ (blue points in Figures 5 and 6), which is not monotonic, and varies considerably between the various samples, is accurately fitted. Moreover, in spite of the strong hypothesis considering an effective lipid tail group in the Matryoshka model, whereas both DMPC and DMPG are known to have two tails, the fits of the amplitude *A*_3_ (red points and lines), including the tail motions, are quite robust among all samples.

Around Q = 1.5 Å^−1^, a little peak can be observed in the experimental points, which is not fitted by the theoretical model. This feature, known as the chain correlation peak (as reported in [53] or [54]), is a Bragg peak caused by the ordering of the alkyl chains, and has a coherent, structural origin. Therefore, it is particularly visible in the d54-MLB, which presents more coherent scattering. However, the current model focuses on the sole incoherent part of the neutron scattering function, thus the local dynamics, and in consequence does not describe the chain correlation peak. In the fit procedure, the points around Q = 1.5 Å^−1^ were then ignored.

The fit parameters, shared between all four amplitudes *A_i_* of one sample at a particular temperature, were retrieved from the fitting procedure, and are displayed in Figure 7 for IN6 measurements, and Figure 8 for IN5 data. The notations are those presented in Section 2.

For every temperature and sample, the retrieved parameters in Figure 7 and 8 exhibit the relevant orders of magnitude known from the literature, however all the distance values in Å are smaller than the structural values. For example, the head radius, *b*_H_, is estimated by diffraction experiments (see [10] and [38]) to be around 4 Å for DMPC (see Figure 2a). However, in Figure 7, the parameter lies within 1 to 2 Å, so more than half less. Similar observations can be done for *R*_H_, which can be estimated in first approximation by 0.5·*d*_H_ sin(*α*). Following diffraction values, 0.5 · *d*_H_ 5 Å, and with *α* = 32.3° (Table 3), we should get a value around 2.6 Å, whereas the *R*_H_ stands around 0.5 and 1 Å in Figure 7. The same effect appears for the tail parameter *R*_1_, which represents the smallest radius in which the lipid tail group is diffusing. It can be directly linked to the H-C-H distance in a methylene and methyl group, and is estimated to be around 2 Å [55], but the values reported here are around 0.5 Å. Partly, this compression of the distances can be explained by a projection effect inherent to the neutron instrument’s setup. The in-plane and out-of-plane directions hold exactly for one detector only, and are more or less mashed with the other direction for all other detectors. The compressed distances are therefore due to a projection representing a mix of the two directions. Moreover, we remind that the parameters obtained are extracted through a dynamical model, so they could be understood more as a deviation from a mean value. They are then better referred as apparent or dynamical values, more than static physical values obtained from diffraction experiments. As a consequence, only the comparison between different samples, or trends with temperature, should be taken into account rather than their absolute value. In general, dynamical and structural values are based on different theories and assumptions, and thus are not directly comparable, as already discussed in [33].

Concerning the other distance parameter, *R*_∥_, accounting for the lateral diffusion radius, the values are also quite small, around 1 - 2 Å. However in that case, one has to consider it in relation with the instrumental resolution of each instrument, ~10 ps for IN6 and ~15 ps for IN5. At such short time scales, the full 2D-diffusion of a lipid within a membrane is not visible, but only the part observable at this resolution. As a consequence, the diffusion radius *R*_∥_ is smaller than the real diffusional free path, that can be of the order of the nm or μm, as can be measured by fluorescence measurements [56].

The force constant *k_force_*, retrieved from the MLBs at 45° (see Figure 7) presents values between 1 and 3 N/m. In elastic neutron scattering studies, an average force constant can be determined with the Bicout-Zaccai model as described in [57], and its application to DMPC membranes leads to values around 1 N/m in the gel phase and 0.2 N/m in the liquid phase [35]. The *k_force_* values displayed here are slightly higher, but they account for the out-of-plane motions. In this direction, the membrane is stiffer as more energy is required for a lipid to move perpendicular to the membrane. Following this consideration, the force constant is higher for out-of-plane motions than the total average in all directions [36].

Regarding the normalization parameters, the mobile fraction *m* is quite high, between 60 and 90 % of all H atoms. We observe an increase with increasing temperature, which is expected as more H atoms will become mobile. Meanwhile, the parameter *ε*_0_ stays between 0 and 30 %, which is smaller than the theoretical immobile fraction, 1 − *m*. However, this error term also accounts for multiple scattering effects, or other experimental errors, as derived in [33].

In parallel, most of the parameters displayed in Figure 7 and 8 have small error bars, except for some values (especially among the head group parameters) which tend towards zero and exhibit high error bars. This behaviour turns up to be purely numerical and caused by the very similar shape of some structure factor functions describing different motions. In particular, the parameters for the head group, *b*_H_ and *R*_H_, appear each in a similar form of the Bessel function (see Table S1 for the detailed expressions), resulting in similar functions for *A_head_* and *A_rot_*. It is then numerically difficult to deconvolute such expressions from the amplitudes *A_i_*. As a consequence, the fitting procedure can lead some parameters to be zero (for example *R*_H_), so that the corresponding amplitude *A_rot_* is equal to one, and does not contribute to the fit.

## 5 Discussion

The effects of the different variations with respect to the samples and instruments are discussed here, then a comparison with existing models is presented.

### 5.1 Impact of temperature and main phase transition

In [36], the same DMPC system was studied by elastic incoherent neutron scattering on IN6 and by neutron diffraction on D16 at ILL, Grenoble. The mean square displacements, consequently the dynamics of the lipid membrane, were shown to increase with temperature, as lipids are gaining more thermal energy. Notably, at the main phase transition temperature *T_m_*, a change of slope appeared. From that, it was shown that *T_m_* is about 296 K for MLVs, and 303 K for MLBs, this discrepancy being largely due to differences in the hydration, the vesicles being more hydrated than bilayers. As a consequence, all the samples at 280 or 283 K are supposed to be in the gel ordered phase, whereas at 311 K and 340 K, they are in the liquid disordered phase. Alternatively, for IN5 data at 295 K, the lipids should be in the middle of the main phase transition.

In general, the increase of motions with temperature is reproduced by the Matryoshka model, noticeable by the rise of the distance parameters. The effect is more striking for the lipid lateral diffusion radius, *R*_∥_, which is almost doubled for the protonated MLBs or MLVs, as seen in Figure 7. On the contrary, the tail parameter *R*_1_ does not display a clear trend, and seems even to tend towards zero at the highest temperature, 340 K, so that the 2D diffusion of the tails is reduced with temperature. It seems counter-intuitive as in the liquid phase the tails are more disorganized. However, in a membrane, the chains are restricted in terms of space, and this could limit, or even prevent, motions when the chains are disorganized and more extended. Such explanation is supported by the *R*_1_ increase for MLVs, where the tails have more space.

The force constant *k_force_* representing the resilience of the membrane in the out-of-plane direction decreases with temperature, passing from high values (3 or 6 N/m) to around 1 N/m in Figure 7. Such decrease indicates an enhanced flexibility, which is consistent with the other parameters, as discussed above, especially after the main phase transition, where the lipids enter the liquid phase.

### 5.2 In-plane against out-of-plane motions

In Figure 7, in-plane and out-of-plane motions are respectively compared through the DMPC MLB135 (blue points) and DMPC MLB45 (orange points). As the Matryoshka model is different for each direction, some parameters change from one direction to another. For example, the lateral diffusion radius *R*_∥_ is only visible in-plane, whereas the force constant *k_force_* only appears for out-of-plane motions. In contrast, the expression for the head rotation and head-flip-flop motions varies with the direction, which could explain the discrepancies in *b*_H_ at 311 K and 340 K.

The probability *ϕ* of jump events tends to be almost constant, about 20 %, when viewed from the normal direction, whereas an abrupt increase from 10 % to 30 % is seen at 340 K for in-plane MLBs. This observation suggests an anisotropy of jump-diffusion in lipids, contrary to the case of methyl groups in proteins. Meanwhile, the MLVs (in green) present a probability *ϕ* close to zero, which is neither close to in-plane nor to out-of-plane motions, although the MLVs represent an average over all directions (four in the current calculations). In parallel, the independent linewidths analysis in [44] shows that for fast motions, the corresponding correlation time *τ*_3_ differs for the in-plane or out-of-plane motions. It supports the anisotropy hypothesis, but we would need more data to fully validate it.

Finally, the mobile fraction *m* tends towards a 5 to 10 % bigger proportion of H atoms involved in out-of-plane displacements than in-plane, even if the amplitude of such motions should be smaller due to the higher energy required to move outside the membrane.

### 5.3 Deuteration of the tails and focus on the head group

Measurements on d54-MLBs in the direction of the membrane enable to focus on the head group, as deuterium atoms in the tails are almost invisible compared to hydrogen atoms in incoherent neutron scattering (see corresponding cross sections of H and D in [26]). Figure 7 displays the comparison of original MLBs (blue points) against d54-MLBs (pink points). Rotation of the whole lipid around its normal axis, represented by the *R*_H_ parameter, seems better determined for deuterated samples, as less parameters are considered in the fits compared to protonated MLBs.

Then, the probability *ϕ* of jump events is much higher for deuterated samples, indicating that jump-diffusion is more likely to happen within the head, compared to the whole lipid (tails included). However, the head group is directly in contact with the hydration layer, whereas the chains are buried within the membrane, and have much less interaction with the water molecules. In consequence, the dynamics in the head groups should be larger, as shown in [35] for mean-square displacements. It is in agreement with what we see with the jump-diffusion, but also the lateral diffusion radius *R*_∥_, which is bigger for deuterated MLBs.

Lastly, the mobile fraction of H atoms is much larger, about 20 % more, for the head group, than for the whole lipid, which is supporting the enhancement of the dynamics of lipid groups in contact with the hydration layer.

In contrast, and in agreement with the hierarchy of motions of the Matryoshka model, the linewidths analysis in [44] reveals that the motions from the head are slower than the whole lipid motions, and thus slower than the tails’ motions. Such results, combined to our observations from the amplitudes’ analysis, indicate that the hydration layer would favor larger exploration (within the membrane, or through jump-diffusion), but higher interactions would slow down the corresponding motions. It shows the importance of distinguishing the geometry of motions (amplitudes’ analysis) and their diffusive properties (linewidth analysis [44]). For instance, larger motions do not mean necessarily faster motions, it is thus necessary to conduct both types of analyses to retrieve a complete dynamical picture of our systems.

### 5.4 Influence of the membrane geometry on the dynamics

Bilayers and vesicles are compared in Figure 7 through the respective blue and green points. On the whole, vesicles are more mobile than bilayers, as could be seen in the lateral diffusion for *R*_∥_, which is by 10 % to 40 % larger for lipids in a vesicle. Similarly, the mobile fraction *m* is more than 10 % higher at 280 K, becoming equal for MLVs and MLBs only in the liquid phase. This effect was also grabbed in elastic neutron scattering on the same system in [36], as well as in the linewidths analysis from [44], and can be explained by a higher hydration for MLVs than MLBs, and the fact that the lipids are less constrained in a vesicle than in a bilayer.

### 5.5 Effect of the instrumental resolution

IN5 and IN6 data, for which the instrumental resolution was similar, respectively 15 and 10 ps, are compared for DMPC around the temperature 280 K in Figure 8. IN5 data are shown in blue empty squares, compared to IN6 data displayed in light blue filled points. The parameters are quite comparable, as expected from the analog resolutions, except for a slight decrease in IN5 data of *b*_H_, linked to the head group rotation and head-flip-flop, as well as the mobile fraction *m*. The time resolution does not allow for so much differences, however, between both experiments, MLBs were more hydrated during IN6 beamtime than IN5 (27 % weight of water on IN6 against 10 % on IN5), and this slight difference is directly visible by the application of the current model, proving its sensitivity to probe changes caused by conditions such as hydration.

### 5.6 Change of lipid composition

In the IN5 experiment, two different lipid compositions were probed : DMPC and DMPG, for which the head group differs. The PG group is known to be lighter : 92 Da, against 104 Da for the PC head group [50]. In Figure 8, DMPC (blue points) is compared against DMPG (yellow points). The head parameters carry the highest error bars, but the most striking effect is seen for *R*_∥_, which indicates that DMPG lipids diffuse more within the membrane than DMPC, which is consistent with the lighter mass of DMPG. Reversely, the mobile fraction *m* is slightly smaller for DMPG than DMPC, which can be put in parallel with the linewidths analysis [44] where the dynamics of DMPG is shown to be slower than DMPC.

On the other hand, whereas the tails are the same for DMPC and DMPG, we observe a smaller *R*_1_ for DMPG than DMPC in the gel phase at 280K. It could indicate that due to the smaller headgroup of DMPG, the lipids are more packed and thus the tails more constrained than for DMPC. On the contrary, at the same temperature, the probability *ϕ* for jump events is much higher for DMPG than DMPC, which is counter-intuitive, and could be due to numerical effects as *ϕ* for DMPC is equal to zero.

### 5.7 Comparison with existing models

In the present study, we compared data obtained from DMPC and DMPG on two different instruments at ILL, IN6 and IN5 with time windows of 10 and 15 ps, respectively. The motions to which this gives access are thus rather short and localized, although at least parts of collective dynamics are also visible and important to include. We are comparing the results to those obtained by Pfeiffer et al. [25] obtained from dipalmitoylphosphatidylcholine (DPPC) and chain deuterated DPPC-*d*_62_ on the spectrometers IN10 and IN5 at ILL, having time windows of about 1 ns and 20 ps, respectively. Further to results from Wanderlingh et al. [30, 31] obtained from DMPC and 1-palmitoyl-oleoyl-*sn*-glycero-phosphocholine (POPC) on IN5 with a 100 ps time window configuration. Finally to results from Gupta and Schneider [24] obtained from four different phospholipid liposome samples, DOPC (1,2-dioleoyl-*sn* − *glycero*-3-phosphocholine), DSPC (1,2-distearoyl-*sn* − *glycero*-3-phosphocholine), DMPC and SoyPC (L-*α*-phosphatidylcholine) on a neutron spin echo (NSE) spectrometer (5 ns < t < 100 ns) and by QENS (t < 5 ns).

In the oldest model of Pfeiffer et al. [25], ordered membranes were measured in in-plane and out-of-plane directions and a rather broad time range covered. The authors included specific motions of the head groups and the tails, in-plane and out-of-plane diffusional motions, rotations along the molecule axis and collective undulations. It included already an exhaustive ensemble of possible local dynamics and was successful in describing the experimental EISF of the samples permitting to extract diffusion constants corresponding to various movements, the distance of protons from rotational axis, the vibrational amplitude and residence times. For long years, it was the model of reference to analyze lipid membrane dynamics. However, only few points in Q were shown, probably due to limited statistics, and no error bars were given for the EISF. It comprised data from two time scales, and first attempts were made to separate the motions according to their typical duration. We can conclude that many details of the Matryoshka are already there, but modern instruments allow to go beyond that model.

The model suggested by Wanderlingh et al. [30, 31] presented clearly that the motions in lipid membranes can be considered as dynamically independent and separated in three time domains within a very good approximation. Such finding allowed to analyze the QENS curves using three Lorentzian functions with widths differing by a factor of about 5 among each other. The authors were able to calculate integrated areas of the Lorentzian functions and to determine EISF of the three motions identified for fast, intermediate and slow dynamics. Fast dynamics were described by rotational diffusion of H atoms with respect to the bounding carbon atom. Intermediate motions were related to lipid chain dynamics; for that the authors subdivided the lipid molecule into beads representing head groups, chain segments and tail methyls. Slow dynamics were given as translational diffusion of the whole phospholipids. Although no chain deuterated lipids were measured, in-plane and out-of-plane motions were invoked to describe the latter one. This model permits to describe rather precisely the EISF and widths of the Lorentzians for different lipid systems and at various temperatures (room temperature and more recently also at 248 and 273 K, corresponding to the lipid gel phase). Still measurements from different orientations and selectively deuterated samples are missing in this study, even if the authors take into account in-plane and out-of-plane motions.

Gupta and Schneider combined QENS and spin-echo techniques to investigate mainly long time dynamics [24]. They also present a combination of independent movements corresponding to different time scales. Fast motions are here described as particles diffusing in a sphere or a cylinder, essentially of the lipid tail, the lipid head group being included as a constant background only. Slower motions, probed by NSE, are modeled by diffusive translational motions, height-height correlations and thickness fluctuations. The three latter movements are seen in time domains not accessible by the spectrometers used in the present study and in this sense the Gupta model is complementary to the Matryoshka model. As all measurements were done with vesicles, no in- or out-of-plane motions could be distinguished. The use of partially deuterated samples led the authors confirm that the head group motions could be mostly ignored.

The development of the Matryoshka model is based on measurements of vesicles and ordered membranes, allowing to compare in-plane and out-of-plane motions and to separate head group and chain motions due to the investigation of partially deuterated samples. It is successfully applied here to results from two different instruments with slightly different resolutions and three different temperatures, corresponding to the gel and fluid phase of the two investigated lipids. To the best of our knowledge, it is therefore the most complete modeling for local motions in lipid vesicles in membranes today.

In addition, attention has been taken to avoid overfitting. First, by fixing some known parameters, based on structural results of many experimental measurements reported in the literature, as the tilt angle of the head *α* or chain length *M*. Then, by performing a global fit of the four amplitudes *A_i_*(*Q*), with shared parameters, which really constrains the search range. Among all the models treating lipid dynamics across time-scales, the Matryoshka model is the first one to apply this method.

Still some parameters must be treated as dynamical values, not in perfect agreement with static results in the literature, what could be due to projections which average over various views and due to the treatment of the two tails as one effective group.

## 6 Conclusion

In this study, we presented a new approach to describe motions in lipid model membranes, ranging from localized ones to collective movements of the whole molecule, as obtained from quasi-elastic neutron scattering data. It is based on well-known molecular dynamics found in lipids and membranes and structural parameters obtained by other methods. Applying a global fit strategy to our data permits to restrain the free parameters further.

The Matryoshka model proves to be successful in describing the data of various types of samples, geometries, temperatures or compositions of lipids. With data fitting of the EISF and Lorentzian amplitudes of good quality, rather small error bars, and reproducibility thorough two experiments from different instruments of similar resolution, it also demonstrates its sensitivity and precision to disentangle subtle differences between samples. These observations, in addition to be supported by literature, form the basis of a more complete dynamical study of standard systems like DMPC or DMPG.

The requirements of much less hypotheses and more robust data fitting with shared parameters are invariably great improvements offered by this new model. With its capacity to probe little variations caused by temperature, hydration or geometry, its range of applicability now awaits to be extended to more complex systems, like other types of lipids or mixtures.

In a forthcoming publication, we will extend further the study to the half-widths at half-maximum of the Lorentzian functions in Equation (1) to validate that the Matryoshka model is also successful in describing the diffusive nature of atomic motions. Later on, a more detailed investigation on collective motions, as studied by inelastic neutron scattering, will also be considered to complement the work of [24]. The variations of atomic motions when crossing the lipidic phase transitions would be worth to be studied in the future.

The fact that all amplitudes are fitted within the Matryoshka model allows to adapt it for data analysis in other contexts of biophysical relevance: recently, A. Cisse et al. [58] used a version inspired from this model to fit data of Apolipoprotein B-100 in interaction with detergent what permitted to separate the dynamical contributions of the two components. The partition *z* is not done between heads and tails here, but between the two components. Such separation is extremely difficult otherwise. The Matryoshka model was also used in a study recently submitted for publication [59] to characterize the structure and dynamics of short chain lipids and alcohols assembled in MLVs. They are mimicking protomembranes at the origin of life and despite the different geometries of the molecules, our model was successful in identifying the parameters key for a correct functionality of the membrane at high temperature.

## Abbreviations

AFM: atomic force microscopy
cryo-EM: cryo-electron microscopy
DMPC: 1,2-dimyristoyl-*sn*-glycero-3-phosphocholine
DMPG: (1,2-dimyristoyl-*sn*-glycero-3-phospho-(1’-rac-glycerol) (sodium salt)
EINS: elastic incoherent neutron scattering
EISF: elastic incoherent structure factor
ILL: Institut Laue Langevin
LAMP: Large Array Manipulation Program
MLBs: multilamellar bilayers
MLVs: multilamellar vesicles
NMR: nuclear magnetic resonance
NSE: neutron spin-echo
QENS: quasi-elastic incoherent neutron scattering
QISF: quasi-elastic incoherent structure factor
SAS: small-angle scattering

## Author contributions

CRediT model (see BBA website : BBA CRediT).

**Aline Cisse :** Software, Validation, Formal Analysis, Writing - Original Draft. **Tatsuhito Matsuo :** Software, Validation, Formal Analysis, Writing - Review & Editing. **Marie Plazanet :** Investigation, Resources. **Francesca Natali :** Conceptualization, Methodology, Investigation. **Michael Marek Koza :** Resources. **Jacques Ollivier :** Resources. **Dominique J. Bicout :** Conceptualization, Methodology, Writing - Review & Editing, Supervision, Funding acquisition. **Judith Peters :** Conceptualization, Methodology, Investigation, Writing - Review & Editing, Supervision, Funding acquisition.

## Acknowledgements

The authors thank the Institut Laue Langevin for beam time to perform the experiment. AC is supported by the Foundation JP Aguilar for her PhD thesis.

## Supplementary Material

## Appendix A Expression of the theoretical EISF

**Table 1:**
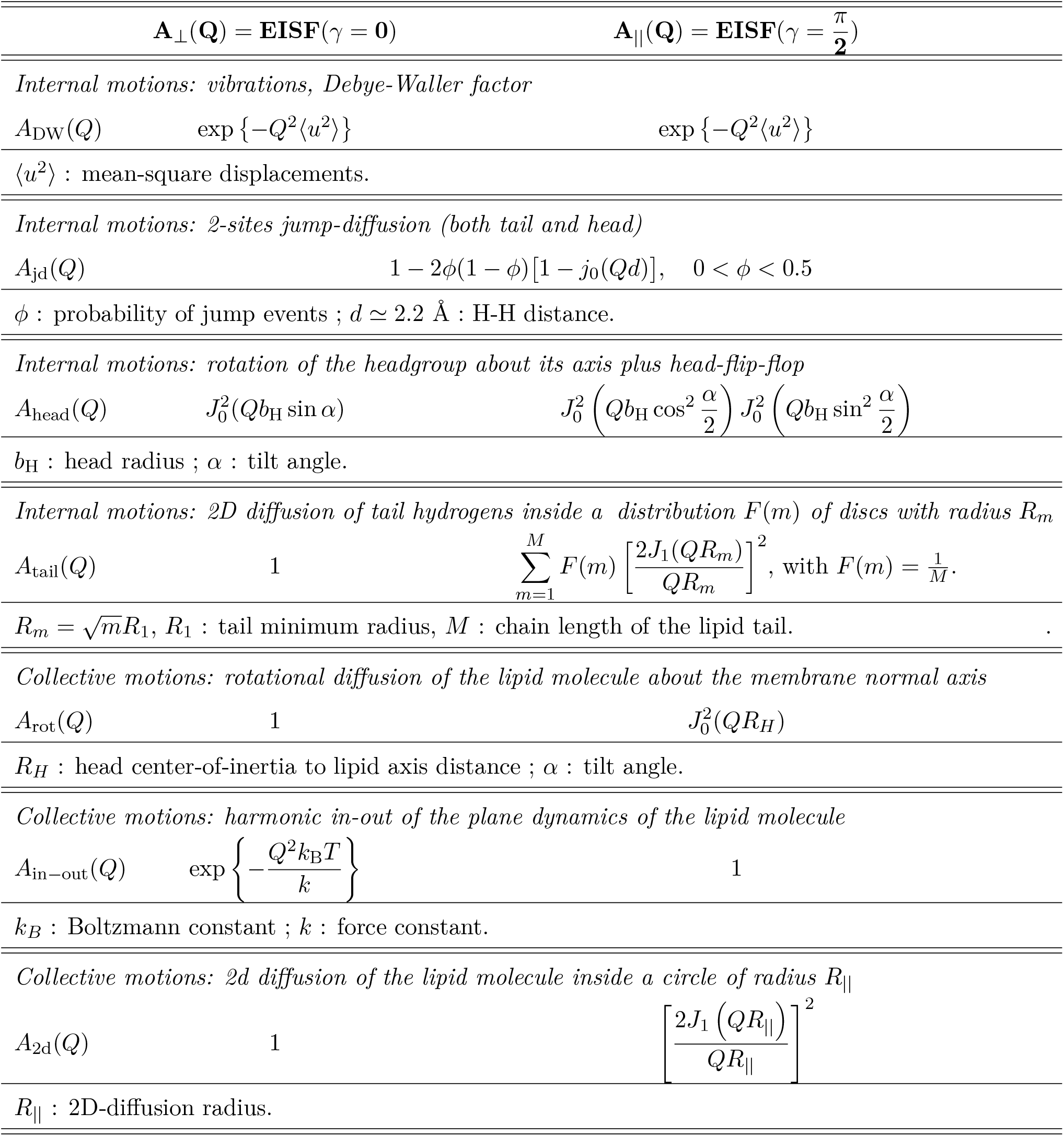
EISF of each motion. The parameters are explicited and can be retrieved in Fig. 2a.

## Appendix B Fit curves against data points for IN6 and IN5 data at higher temperatures

**Figure 1:**
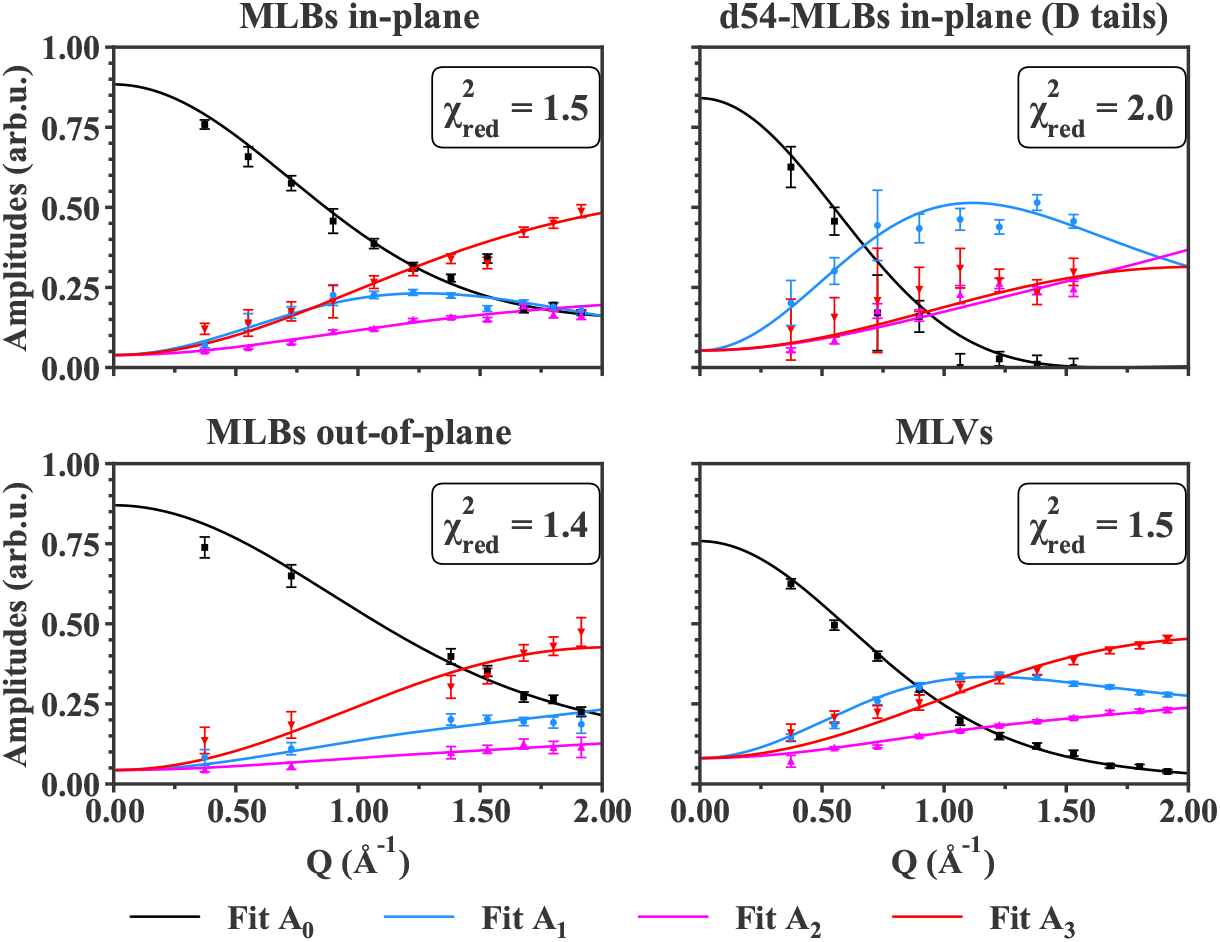
Fit curves against data points for IN6 data at T = 311K.

**Figure 2:**
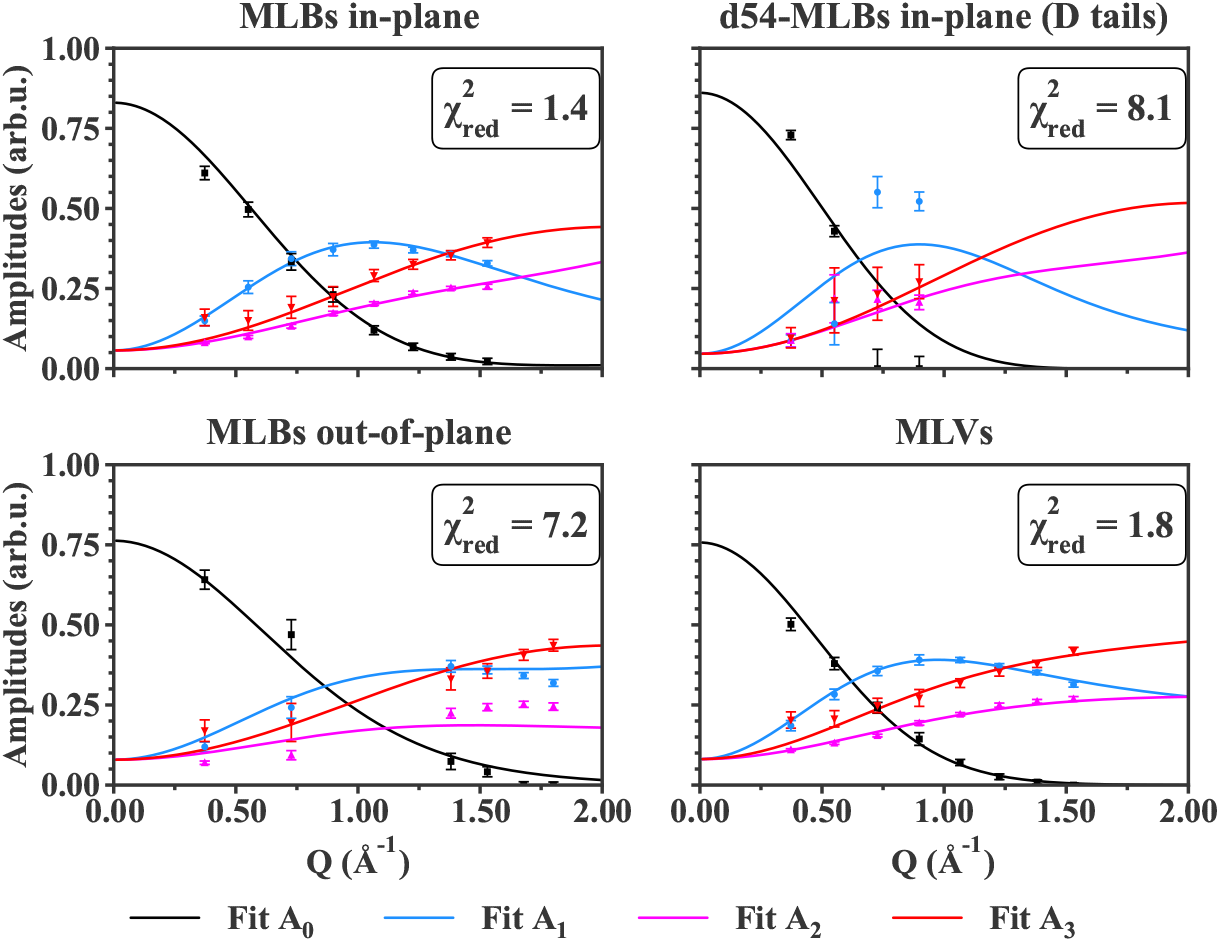
Fit curves against data points for IN6 data at T = 340K.

**Figure 3:**
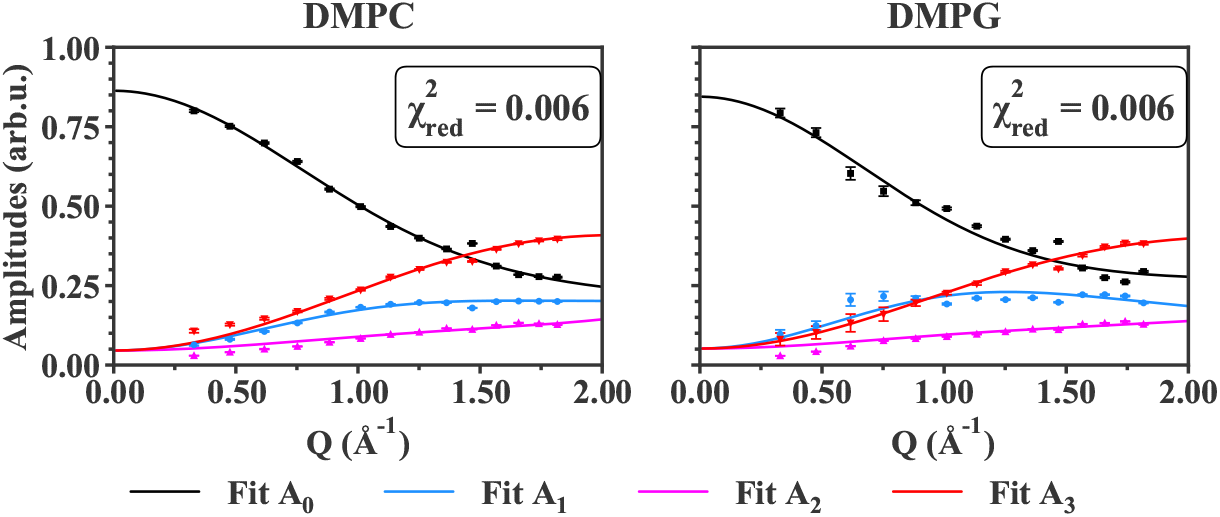
Fit curves against data points for IN5 data at T = 295K.

## Appendix C Comparison between samples of the fit curves

**Figure 4:**
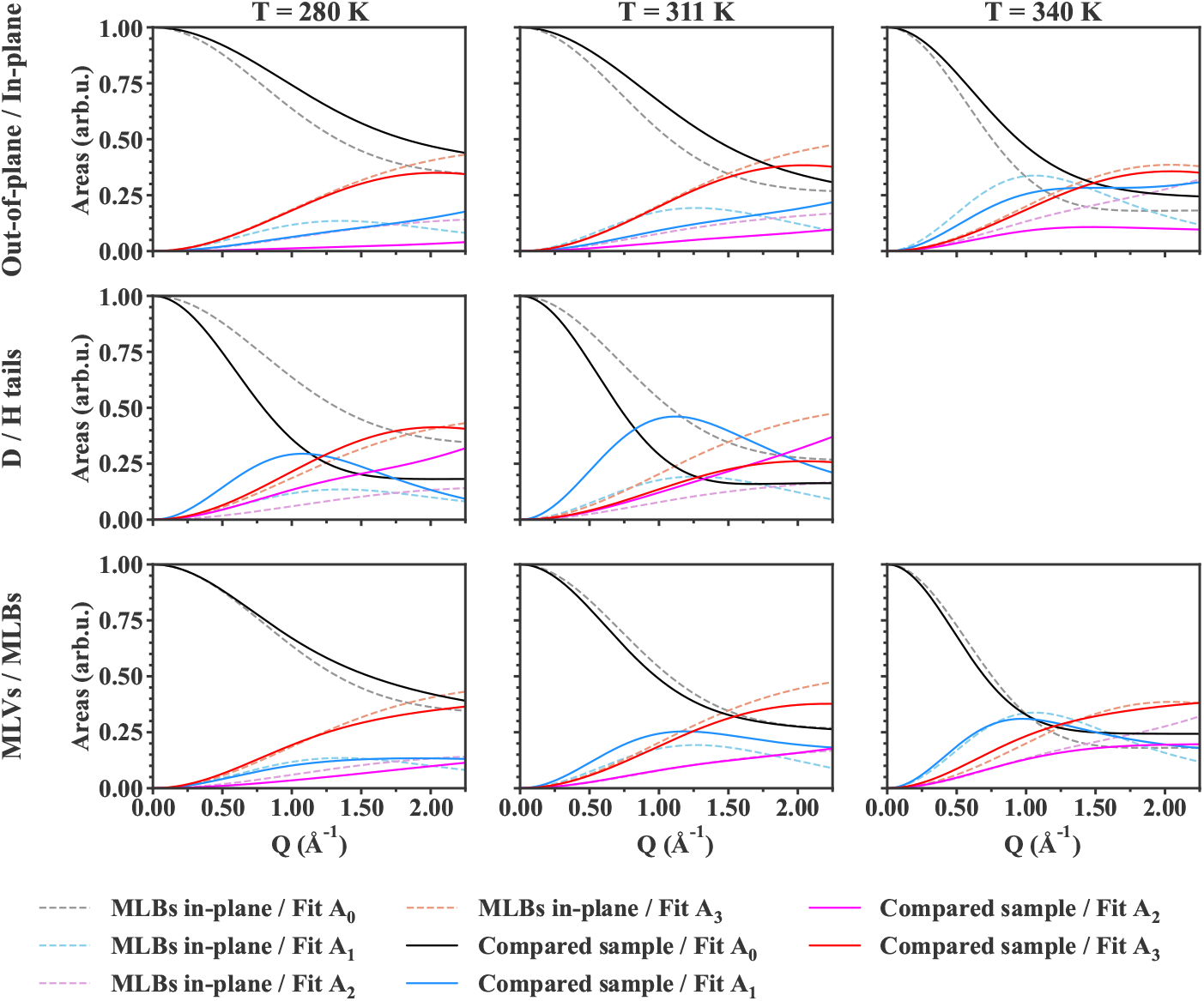
Comparison of the fits for IN6 samples.

**Figure 5:**
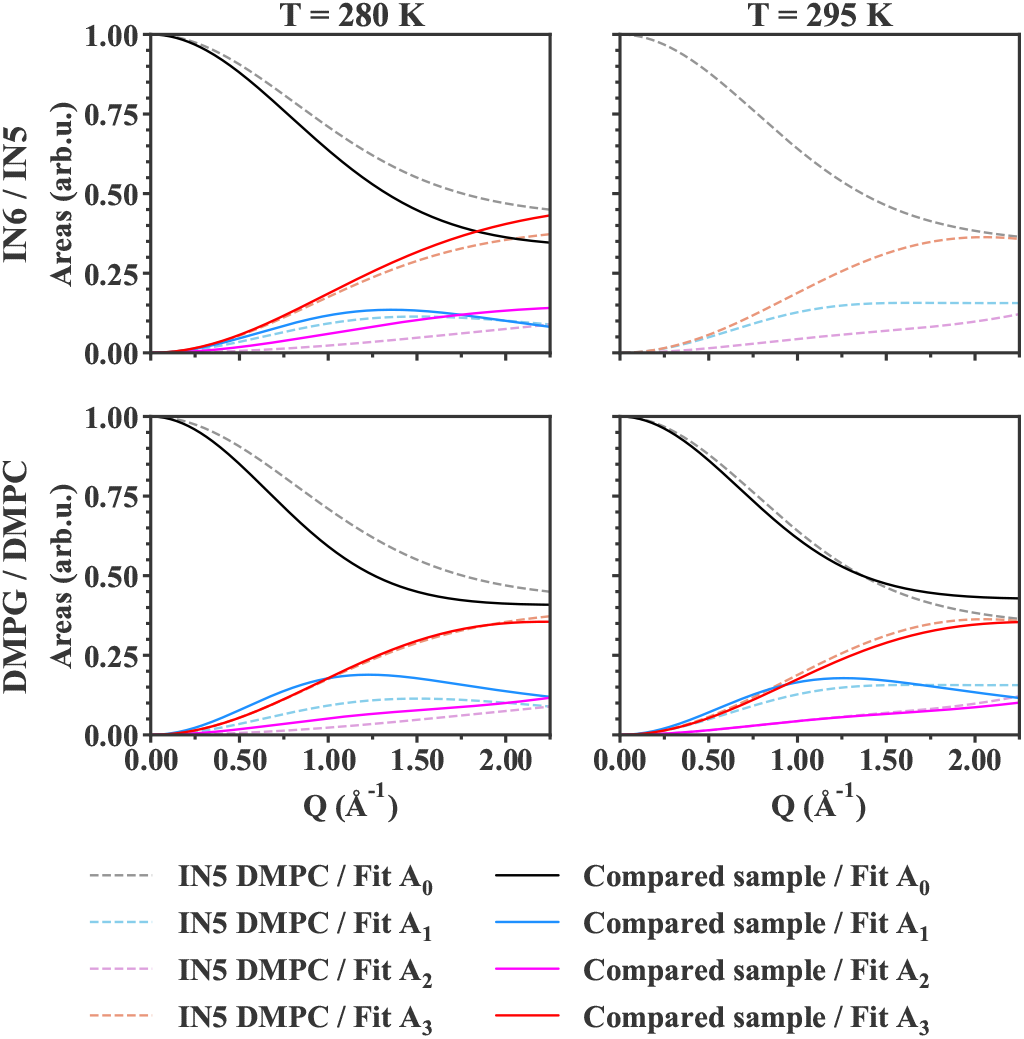
Comparison of the fits for IN5 samples.

## Appendix D Table of results

### D.1 IN6 : DMPC MLBs measured at 135° (in-plane)

**Table 2:** Fit parameters and its corresponding standard error (95% confidence interval) for DMPC MLBs sample measured on IN6 at 135° (in-plane motions). 0.01 is the minimum boundary in the fit (that is why values are not equal to 0 but 0.01).

| Parameter | T = 280 K | T = 311 K | T = 340 K |
| --- | --- | --- | --- |
| $b_H$ (Å) | $1.04 \pm 0.02$ | $0.62 \pm 0.02$ | $0.66 \pm 0.03$ |
| $R_H$ (Å) | $0.01 \pm 0.42$ | $0.46 \pm 0.01$ | $0.62 \pm 0.01$ |
| $R_1$ (Å) | $0.43 \pm 0.01$ | $0.42 \pm 0.01$ | $0.01 \pm 0.2$ |
| $\phi$ | $0.06 \pm 0.01$ | $0.07 \pm 0.01$ | $0.26 \pm 0.01$ |
| $R$ (Å) | $1.13 \pm 0.03$ | $1.38 \pm 0.02$ | $2.06 \pm 0.03$ |
| $k_{\text{force}}$ (N/m) | / | / | / |
| $m$ | $0.67 \pm 0.01$ | $0.74 \pm 0.02$ | $0.82 \pm 0.01$ |
| $\varepsilon_0$ | $0.24 \pm 0.01$ | $0.15 \pm 0.01$ | $0.01 \pm 0.01$ |

### D.2 IN6 : DMPC MLBs measured at 45° (out-of-plane)

**Table 3:**
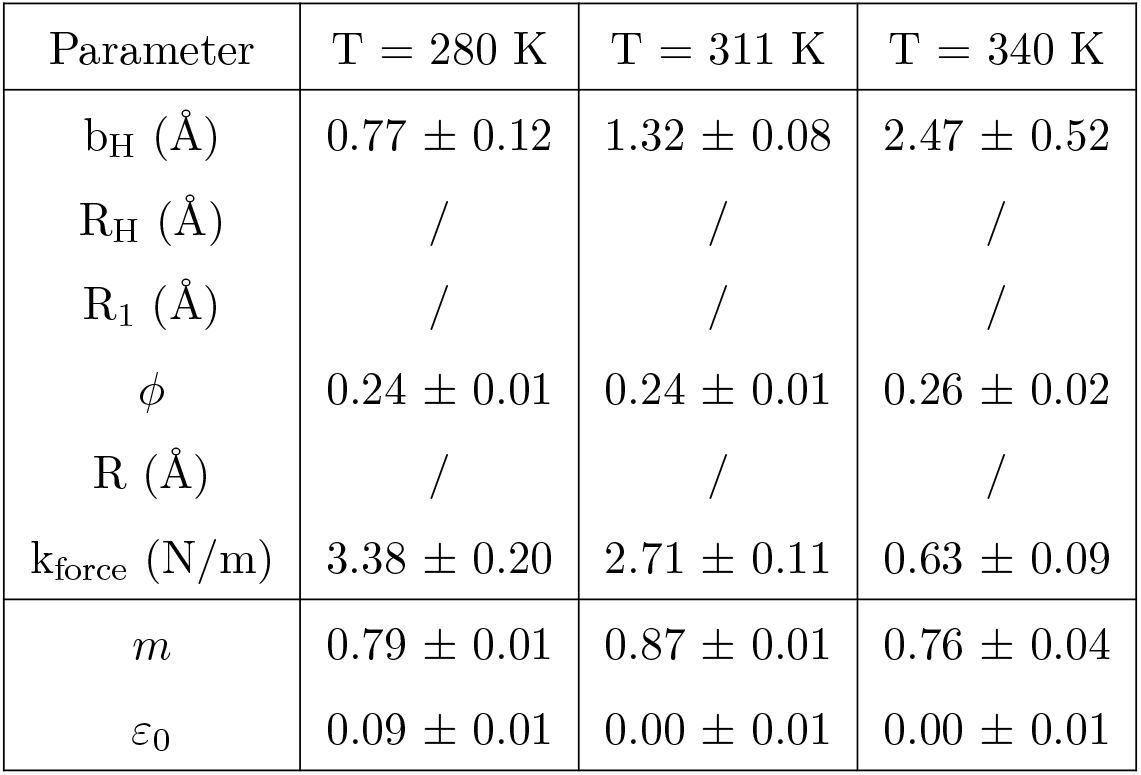
Fit parameters and its corresponding standard error (95% confidence interval) for DMPC MLBs sample measured on IN6 at 45° (out-of-plane motions). 0.01 is the minimum boundary in the fit (that is why values are not equal to 0 but 0.01).

| Parameter | T = 280 K | T = 311 K | T = 340 K |
| --- | --- | --- | --- |
| $b_H$ (Å) | $0.77 \pm 0.12$ | $1.32 \pm 0.08$ | $2.47 \pm 0.52$ |
| $R_H$ (Å) | / | / | / |
| $R_1$ (Å) | / | / | / |
| $\phi$ | $0.24 \pm 0.01$ | $0.24 \pm 0.01$ | $0.26 \pm 0.02$ |
| $R$ (Å) | / | / | / |
| $k_{\text{force}}$ (N/m) | $3.38 \pm 0.20$ | $2.71 \pm 0.11$ | $0.63 \pm 0.09$ |
| $m$ | $0.79 \pm 0.01$ | $0.87 \pm 0.01$ | $0.76 \pm 0.04$ |
| $\varepsilon_0$ | $0.09 \pm 0.01$ | $0.00 \pm 0.01$ | $0.00 \pm 0.01$ |

### D.3 IN6 : d54-DMPC MLBs measured at 135° (in-plane)

**Table 4:** Fit parameters and its corresponding standard error (95% confidence interval) for d54-DMPC MLBs sample measured on IN6 at 135° (in-plane motions). 0.01 is the minimum boundary in the fit (that is why values are not equal to 0 but 0.01).

| Parameter | T = 280 K | T = 311 K | T = 340 K |
| --- | --- | --- | --- |
| $b_H$ (Å) | $0.01 \pm 1.84$ | $0.47 \pm 0.03$ | $0.66 \pm 0.23$ |
| $R_H$ (Å) | $0.70 \pm 0.02$ | $0.43 \pm 0.03$ | $0.61 \pm 0.21$ |
| $R_1$ (Å) | / | / | / |
| $\phi$ | $0.29 \pm 0.03$ | $0.15 \pm 0.02$ | $0.34 \pm 0.32$ |
| $R$ (Å) | $1.89 \pm 0.08$ | $2.28 \pm 0.09$ | $2.35 \pm 0.19$ |
| $k_{\text{force}}$ (N/m) | / | / | / |
| $m$ | $0.82 \pm 0.02$ | $0.84 \pm 0.03$ | $0.86 \pm 0.05$ |
| $\varepsilon_0$ | $0.15 \pm 0.01$ | $0.00 \pm 0.01$ | $0.00 \pm 0.01$ |

### D.4 IN6 : DMPC MLVs measured at 135°

**Table 5:** Fit parameters and its corresponding standard error (95% confidence interval) for DMPC MLVs sample measured on IN6 at 135°. 0.01 is the minimum boundary in the fit (that is why values are not equal to 0 but 0.01).

| Parameter | T = 280 K | T = 311 K | T = 340 K |
| --- | --- | --- | --- |
| $b_H$ (Å) | $0.70 \pm 0.02$ | $0.53 \pm 0.04$ | $1.69 \pm 0.03$ |
| $R_H$ (Å) | $0.29 \pm 0.02$ | $0.74 \pm 0.01$ | $0.58 \pm 0.04$ |
| $R_1$ (Å) | $0.73 \pm 0.01$ | $0.20 \pm 0.01$ | $0.87 \pm 0.02$ |
| $\phi$ | $0.00 \pm 0.01$ | $0.23 \pm 0.01$ | $0.00 \pm 0.01$ |
| $R$ (Å) | $1.33 \pm 0.03$ | $2.15 \pm 0.05$ | $1.98 \pm 0.11$ |
| $k_{\text{force}}$ (N/m) | $5.78 \pm 0.43$ | $1.25 \pm 0.05$ | $0.26 \pm 0.01$ |
| $m$ | $0.77 \pm 0.01$ | $0.76 \pm 0.01$ | $0.76 \pm 0.01$ |
| $\varepsilon_0$ | $0.05 \pm 0.01$ | $0.00 \pm 0.01$ | $0.00 \pm 0.01$ |

### D.5 IN5 : DMPC MLBs measured at 135° (in-plane)

**Table 6:** Fit parameters and its corresponding standard error (95% confidence interval) for DMPC MLBs sample measured on IN5 at 135° (in-plane motions). 0.01 is the minimum boundary in the fit (that is why values are not equal to 0 but 0.01).

| Parameter | T = 280 K | T = 295 K |
| --- | --- | --- |
| $b_H$ (Å) | $0.63 \pm 0.18$ | $0.01 \pm 2.79$ |
| $R_H$ (Å) | $0.01 \pm 0.04$ | $0.44 \pm 0.07$ |
| $R_1$ (Å) | $0.54 \pm 0.04$ | $0.04 \pm 0.24$ |
| $\phi$ | $0.00 \pm 0.04$ | $0.34 \pm 0.05$ |
| $R$ (Å) | $1.05 \pm 0.11$ | $1.18 \pm 0.09$ |
| $k_{\text{force}}$ (N/m) | / | / |
| $m$ | $0.57 \pm 0.03$ | $0.66 \pm 0.03$ |
| $\varepsilon_0$ | $0.29 \pm 0.02$ | $0.21 \pm 0.02$ |

### D.6 IN5 : DMPG MLBs measured at 135° (in-plane)

**Table 7:**
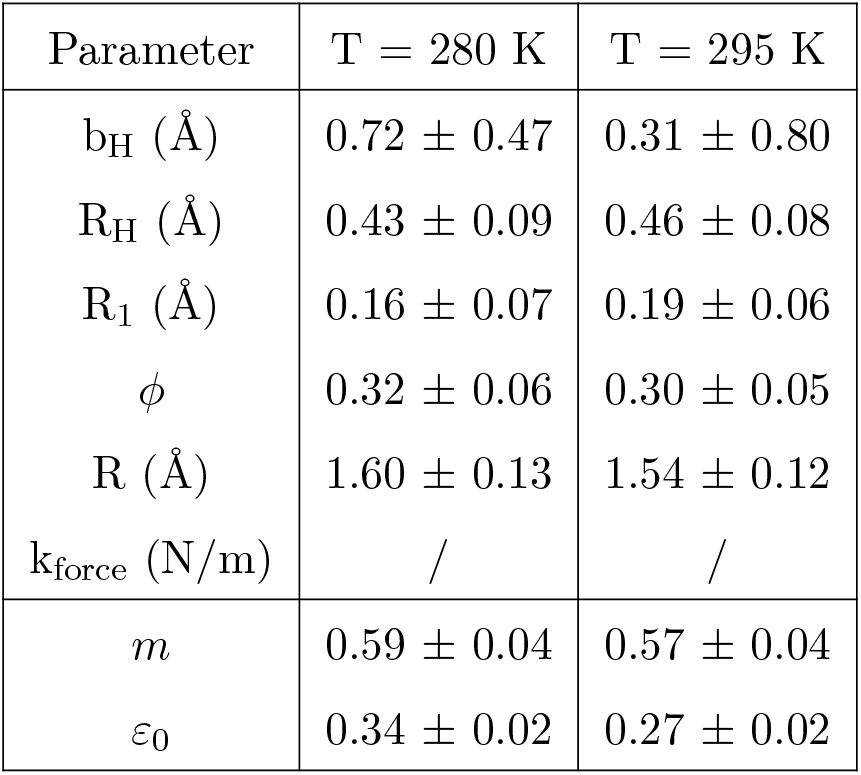
Fit parameters and its corresponding standard error (95% confidence interval) for DMPG MLBs sample measured on IN5 at 135° (in-plane motions). 0.01 is the minimum boundary in the fit (that is why values are not equal to 0 but 0.01).

## Appendix E Other version of the model that does not fit the data

**Table 8:** Other fit functions tested for the areas. *p_imm_* refers to the immobile fraction of Hydrogen atoms. *a* is a factor accounting for the mobile fraction of H atoms, but also for multiple scattering effects that are visible at low-Q range.

| Areas | Theoretical function |
| --- | --- |
| $A_0$ (EISF) | $p_{\text{imm}} + a \left[ [(1-z)A_{\text{tail}} + zA_{\text{head}}] A_{\text{rot}} A_{\text{in-out}} A_{2d} \right]$ |
| $A_1$ (slow) | $\frac{1-a-p_{\text{imm}}}{3} + a \left[ [(1-z)A_{\text{tail}} + zA_{\text{head}}] (1 - A_{\text{rot}} A_{\text{in-out}} A_{2d}) \right]$ |
| $A_2$ (intermediate) | $\frac{1-a-p_{\text{imm}}}{3} + a \left[ [1 - ((1-z)A_{\text{tail}} + zA_{\text{head}})] (A_{\text{rot}} A_{\text{in-out}} A_{2d}) \right]$ |
| $A_3$ (fast) | $\frac{1-a-p_{\text{imm}}}{3} + a \left[ [1 - ((1-z)A_{\text{tail}} + zA_{\text{head}})] (1 - A_{\text{rot}} A_{\text{in-out}} A_{2d}) \right]$ |

**Figure 6:**
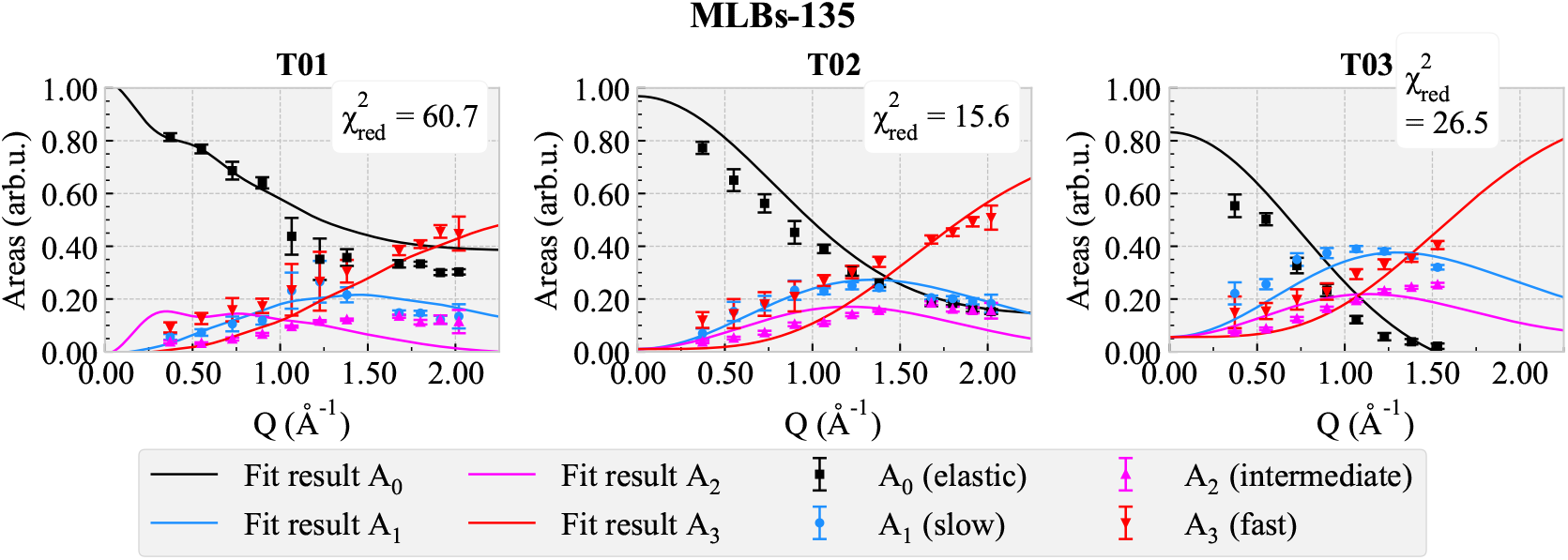
Fit results with the functions gathered in Table 8.

Fig. 6 shows that the fit curves do not match the data points, as well as unstabilities at T = 280K. Moreover, the reduced *χ*^2^ are much higher than for the current model that fits well the four areas (see Fig. 5 for comparison).

## Notes

### Competing Interest Statement

The authors have declared no competing interest.

